# Language networks in aphasia and health: a 1000 participant Activation Likelihood Estimate analysis

**DOI:** 10.1101/2020.06.30.179655

**Authors:** James D. Stefaniak, Reem S. W. Alyahya, Matthew A. Lambon Ralph

**Affiliations:** Division of Neuroscience and Experimental Psychology, University of Manchester, Oxford Road, Manchester, UK; MRC Cognition and Brain Sciences Unit, University of Cambridge, Cambridge, UK; King Fahad Medical City, Riyadh, Saudi Arabia

**Keywords:** Stroke: imaging, Stroke: rehabilitation, Aphasia, Cortical plasticity, Cognitive control

## Abstract

Aphasia recovery post-stroke is classically and most commonly hypothesised to rely on regions that were not involved in language premorbidly, through ‘neurocomputational invasion’ or engagement of ‘quiescent homologues’. Contemporary accounts have suggested, instead, that recovery might be supported by under-utilised areas of the premorbid language network, which are downregulated in health to save neural resources (‘variable neurodisplacement’). Despite the importance of understanding the neural bases of language recovery clinically and theoretically, there is no consensus as to which specific regions are activated more consistently in post-stroke aphasia (PSA) than healthy individuals. Accordingly, we performed an Activation Likelihood Estimation analysis of language functional neuroimaging studies in PSA and linked control data. We obtained coordinate-based functional neuroimaging data for 481 individuals with aphasia following left hemisphere stroke (one third of which was previously unpublished) and for 530 healthy controls. Instead of the language network expanding by activating novel right hemisphere regions ‘de novo’ post-stroke, as would be predicted by neurocomputational invasion/quiescent homologue engagement mechanisms of recovery, we found that multiple regions throughout both hemispheres were consistently activated during language tasks in PSA and controls. Multiple undamaged regions were less consistently activated in PSA than controls, including domain-general regions of medial superior frontal cortex and right fronto-temporal cortex. In the reverse direction, the right anterior insula and inferior frontal gyrus were more consistently activated in PSA than controls, particularly for executively-demanding comprehension tasks. These regions overlap with control networks known to be recruited during difficult tasks in healthy individuals and were more consistently activated by patients during higher than lower demand tasks in this meta-analysis. Overall, these findings run counter to neurocomputational invasion of the language network into new territory or engagement of quiescent homologues. Instead, many parts of the pre-existing language network are less consistently activated in PSA, except for more consistent use of spare capacity within right hemisphere executive-control related regions (cf. variable neurodisplacement). This study provides novel insights into the language network changes that occur post-stroke. Such knowledge is essential if we are to design neurobiologically-informed therapeutic interventions to facilitate language recovery.

## Introduction

Post-stroke aphasia (PSA) is prevalent and debilitating (Engelter *et al*., 2006) and recovery of function tends to be variable and often incomplete (Yagata *et al*., 2017). Compensatory changes in patterns of neural activity, reflecting increased utilisation of surviving neural regions, are hypothesised to contribute to aphasia recovery (Murphy and Corbett, 2009; Turkeltaub *et al*., 2011; Stefaniak *et al*., 2020). While previous studies have explored which set of regions are activated in PSA (Turkeltaub *et al*., 2011), multiple key questions remain unanswered. These include: (a) which specific regions are activated more consistently in PSA than healthy individuals; (b) whether the activated regions differ across different language tasks, and (c) at different stages of recovery. Such knowledge will be essential to understand the mechanisms underlying language network plasticity and thus design neurobiologically-informed therapeutic interventions to aid language recovery. Accordingly, this study tackled these targeted questions through the largest Activation Likelihood Estimation (ALE) analysis, to date, of functional neuroimaging studies in PSA (n=481) and healthy controls (n=530). There were several specific questions we sought to address. We consider these briefly, below, with respect to three major themes.

First, even though recovery of language after stroke has perplexed researchers since the seminal studies of aphasia in the nineteenth century (Finger *et al*., 2003), there have been very few formal, implemented models (Stefaniak *et al*., 2020) and hypotheses have rarely been tested in relation to large patient datasets. Certain mechanisms underlying partial language recovery in PSA propose that neural networks unused in health can adapt after stroke to perform a similar function to the one normally supported by the now damaged neural network(s) (Stefaniak *et al*., 2020), for instance through immediate engagement of *quiescent homologues* (Finger *et al*., 2003) or through *neurocomputational invasion* of non-language regions via experience-dependent plasticity (Keidel *et al*., 2010; Southwell *et al*., 2016). Alternatively, *variable neurodisplacement* (Binney and Lambon Ralph, 2015; Stefaniak *et al*., 2020) proposes that ‘well engineered’ language and cognitive networks dynamically balance performance demand against energy expenditure, downregulating spare capacity under standard performance demands in health but running the remaining system ‘harder’ after partial damage (as the intact system can do when under increased performance demands (Sharp *et al*., 2010; Robson *et al*., 2014; Jung and Lambon Ralph, 2016; Rice *et al*., 2018)). These mechanisms are not mutually exclusive and might include both language-specific and non-language networks, including domain-general executive networks, in both hemispheres (Stefaniak *et al*., 2020). Key predictions of *variable neurodisplacement* are that positive language network changes in PSA are due to upregulation of spare capacity within the pre-existing language network, and that these same upregulated neural regions show increased activation for hard over easier tasks in both PSA and healthy individuals.

Second, there is a tendency to treat ‘language’ and its recovery as a single, homogenous cognitive function. Instead, language refers to a diverse range of expressive and receptive activities. Different language activities are supported by interactions between various more general neurocognitive computations (Patterson and Lambon Ralph, 1999; Gordon *et al*., 2002; Mementi *et al*., 2011) which can be damaged independently of each other to generate the graded, multidimensional nature of post-stroke aphasia (Kummerer *et al*., 2013; Butler *et al*., 2014; Mirman *et al*., 2015; Halai *et al*., 2017; Alyahya *et al*., 2020). Consequently, theories of recovery need to consider not only how each primary neurocognitive system might recover, but also how changes in their interactivity can support improved performance across different language activities. Changes in the division of labour across systems can occur not only between language networks (Ueno *et al*., 2011) but also between language and multi-demand executive systems (Geranmayeh *et al*., 2017; Hartwigsen, 2018).

An important second aspect of this issue is that different subcomponents of language, such as those subserving comprehension versus production, might have differently distributed networks, including degrees of lateralisation, premorbidly (Lidzba *et al*., 2011). For instance, the language network is often described as unilateral (Mazoyer *et al*., 2014) but several lines of evidence suggest it is at least partially bilateral but asymmetric (Lambon Ralph *et al*., 2001; Fedorenko *et al*., 2011). This has significant implications as many studies have highlighted a role for the right hemisphere in recovery (Crinion and Price, 2005; Skipper-Kallal *et al*., 2017a, b). Depending on the degree of premorbid asymmetry, right hemisphere activation might reflect engagement of pre-existing right hemispheric regions of the language network via *variable neurodisplacement* versus novel recruitment of non-language regions via *neurocomputational invasion* (Warburton *et al*., 1999). It is important, therefore, to compare activation patterns in post-stroke aphasia with the natural distribution of the same language subcomponent(s) in healthy individuals.

Third, language recovery is dynamic and occurs most rapidly during the first few months post-stroke (Pedersen *et al*., 1995; Yagata *et al*., 2017), with spontaneous language changes being slower and smaller by the ‘chronic’ stage after approximately 6-12 months (Hope *et al*., 2017). Thus, in order to identify language network changes that are associated with recovery, it is important to compare language networks at subacute vs. chronic stages of recovery.

Given these many outstanding questions, this study sought to identify whether consistent patterns of language-related activation differ between PSA and healthy controls. The omnibus ALE analysis considered which specific regions are activated more consistently in PSA than healthy individuals across all language tasks. Subsequent analyses investigated differences based on: comprehension versus production tasks; for each task type, higher versus lower demand tasks; and time post stroke (e.g., sub-acute vs chronic PSA). If language recovery reflects *neurocomputational invasion* or engagement of *quiescent homologues* then the post-stroke language network should expand to include novel regions that are not consistently activated in healthy individuals, even under increased task difficulty. Conversely, *variable neurodisplacement* predicts that the networks observed in PSA should also be observed in healthy controls, particularly when the healthy system is placed under greater performance demands.

## Materials and methods

### Study search and selection

We searched the databases Medline, Embase and PsycINFO up to April 2020. Terms relating to aphasia (aphasia OR dysphasia OR language OR fluency OR phonology OR semantics OR naming OR repetition OR comprehension OR speaking), stroke (stroke OR ischaemia OR ischemia OR infarct) and neuroimaging (fMRI OR PET OR neuroimaging OR imaging OR functional) were used. We identified eligible articles reporting observational studies that had: a) more than one person with language impairment at any time following left hemispheric stroke; b) more than one healthy control; and c) performed BOLD fMRI or ^15^O-PET during language task-based functional neuroimaging. We extracted coordinate data for inclusion in this ALE meta-analysis that: related to activation (not deactivation) during a language task-based functional neuroimaging experiment; was provided in standard space; was derived from whole-brain mass-univariate analyses without region of interests (ROIs), small volume corrections (SVC), or conjunctions (Müller *et al*., 2018); and was calculated using the same significance thresholds in the PSA and control groups. If coordinates meeting these criteria for both the PSA and control groups were not provided in the publication, the authors were contacted to request unpublished coordinates. Full details are reported in the Supplementary Material.

### ALE meta-analysis

Peak coordinates pertaining to language activation were extracted from each included article and double checked by the same author (JDS). Coordinates in Talairach space were converted to Montreal Neurological Institute (MNI) space using the Lancaster transformation (Lancaster *et al*., 2007). GingerALE 3.0.2 was used to perform ALE (http://brainmap.org/ale/), which is a random-effects coordinate-based meta-analytic technique that identifies neural regions at which activation peaks converge above-chance across studies (Eickhoff *et al*., 2009; Eickhoff *et al*., 2011; Eickhoff *et al*., 2012; Turkeltaub *et al*., 2012). Briefly, we grouped together activation peaks from all imaging tasks performed by the same participant group. Each peak was modelled as a 3D Gaussian distribution of activation probability; each voxel in the brain was assigned the activation probability from the peak within the shortest Euclidean distance, producing a Modelled Activation map for each subject group (Turkeltaub *et al*., 2012). The voxel-wise union of all Modelled Activation maps from all subject groups included in a single dataset produced an ALE map, representing the voxel-wise probability that activation was found for at least one subject group included in that dataset (Turkeltaub *et al*., 2012). Each dataset’s ALE map was thresholded with a voxel-wise uncorrected p<0.001 cluster-forming threshold and a cluster-wise family-wise error (FWE) corrected threshold of p<0.05 based on 1000 random permutations (Eickhoff *et al*., 2016). Coordinates from tasks at different timepoints on the same subject group were not pooled; only tasks performed at the longest timepoint post-stroke for each group were included. If coordinates were available for separate groups within the same study (e.g., for stroke survivors with aphasia as individuals or sub-groups), each individual/sub-group was counted as being from a separate subject group in the meta-analysis. Conjunction images identifying regions in which two datasets both showed convergent activation were created from their voxel-wise minimum thresholded ALE values. Contrast analyses were then performed to identify regions where activation probability significantly differed between two datasets. Thresholded ALE maps from the two datasets being contrasted were subtracted from each other and thresholded at p<0.05 (uncorrected) using 10,000 P-value permutations with a minimum cluster threshold of 200mm^3^. The Harvard-Oxford atlas (Desikan *et al*., 2006) defined anatomical labels and the Talairach Daemon atlas (Lancaster *et al*., 2000) determined the Brodmann Area label associated with each peak coordinate.

We performed a set of pre-planned ALE analyses that are set out below. We required single datasets to have at least 17 subject groups for inclusion in the omnibus ALE meta-analysis, as recommended by empirical simulations suggesting this number was needed to ensure adequate power (Eickhoff *et al*., 2016). Given the scarcity of functional neuroimaging studies in PSA, we required 10 subject groups for single datasets to be included in subgroup ALE analyses for more specific contrasts.

### Omnibus ALE analysis

This analysis combined all data available. Thus, it consisted of single dataset, conjunction and contrast ALE meta-analyses comparing all language tasks in all PSA against all language tasks in all controls.

### Subgroup ALE meta-analyses

Following the omnibus ALE analysis, PSA groups were divided into one of two categories according to a characteristic of the language task performed or a characteristic of its participants. Single dataset, conjunction and contrast ALE meta-analyses were conducted to compare the resultant two categories of PSA groups. Then, each control group was categorised according to how its corresponding PSA group had been categorised and the same ALE meta-analyses were performed on the resultant two categories of controls. Finally, conjunction and contrast ALE meta-analyses were conducted to compare the corresponding categories of PSA to controls. If the same PSA group performed multiple imaging tasks which were divided into different categories, the coordinates for both imaging tasks were included in their respective categories. The following subgroup ALE meta-analyses were conducted:

#### A) Comprehension vs. production

PSA participants might activate different neural regions relative to controls for a subset of language tasks. Such differences may have been obscured by including all language tasks in the omnibus ALE analysis. Functional neuroimaging tasks were therefore categorised according to whether they involved ‘production’ (including either overt or covert production of sublexical, lexical or sentence level speech components) or solely ‘comprehension’ without production.

#### B) Higher versus lower processing demand

*Variable neurodisplacement* proposes that neural spare capacity is downregulated to save energy under standard performance demands in health but is upregulated when performance demands increase post-stroke. If this occurs, we would expect the neural regions upregulated in PSA to be more consistently activated during more difficult compared to less difficult tasks in both PSA and controls. Therefore, comprehension and production tasks were each subdivided according to task difficulty. Higher demand comprehension tasks were defined as tasks requiring a linguistic decision to be made; e.g., whether a stimulus is a word or pseudoword, concrete or abstract, or related to some other semantic or syntactic property. Lower demand comprehension tasks either did not require a linguistic decision or required a very simple identity match; e.g., passive listening or simple word-picture matching. Higher demand production tasks required production of >1 word, such as propositional speech or category fluency tasks. Lower demand production tasks required production of single words, such as picture naming or single item repetition.

#### C) Time post-stroke

Language recovery occurs most rapidly during the first six months post-stroke (Pedersen *et al*., 1995; Yagata *et al*., 2017). PSA groups were therefore categorised according to whether their mean time post-stroke was before or after 6 months.

### Statistical analysis

We compared mean ages of the PSA and control groups using Mann-Whitney U tests implemented in SPSS version 25 with statistical significance defined as p<0.05 with Bonferroni correction.

### Data availability

The data supporting the findings of this study are available within the Supplementary Material.

## Results

### Descriptive statistics

10,169 unique references were obtained from the search. 79 papers were eligible for inclusion; useable foci were obtained from 33/79 included papers. A flowchart of the search and selection process is shown in Fig.1. Details of the included/excluded papers, reasons for excluding eligible papers, and information on the PSA groups included in the ALE analysis are provided in Supplementary Tables S1-3. Across all language tasks, 1521 foci were obtained from 481 PSA in 64 groups, and 809 foci were obtained from 530 healthy controls in 37 groups (Supplementary Tables S3, 4). Foci relating to 172 of the 481 PSA had not been published but were provided after personal communication with the corresponding authors (Schofield *et al*., 2012; Geranmayeh *et al*., 2016; Radman *et al*., 2016; Hallam *et al*., 2018; Wilson *et al*., 2018; Barbieri *et al*., 2019; Meier *et al*., 2019; Tao and Rapp, 2019).

**Figure 1:**
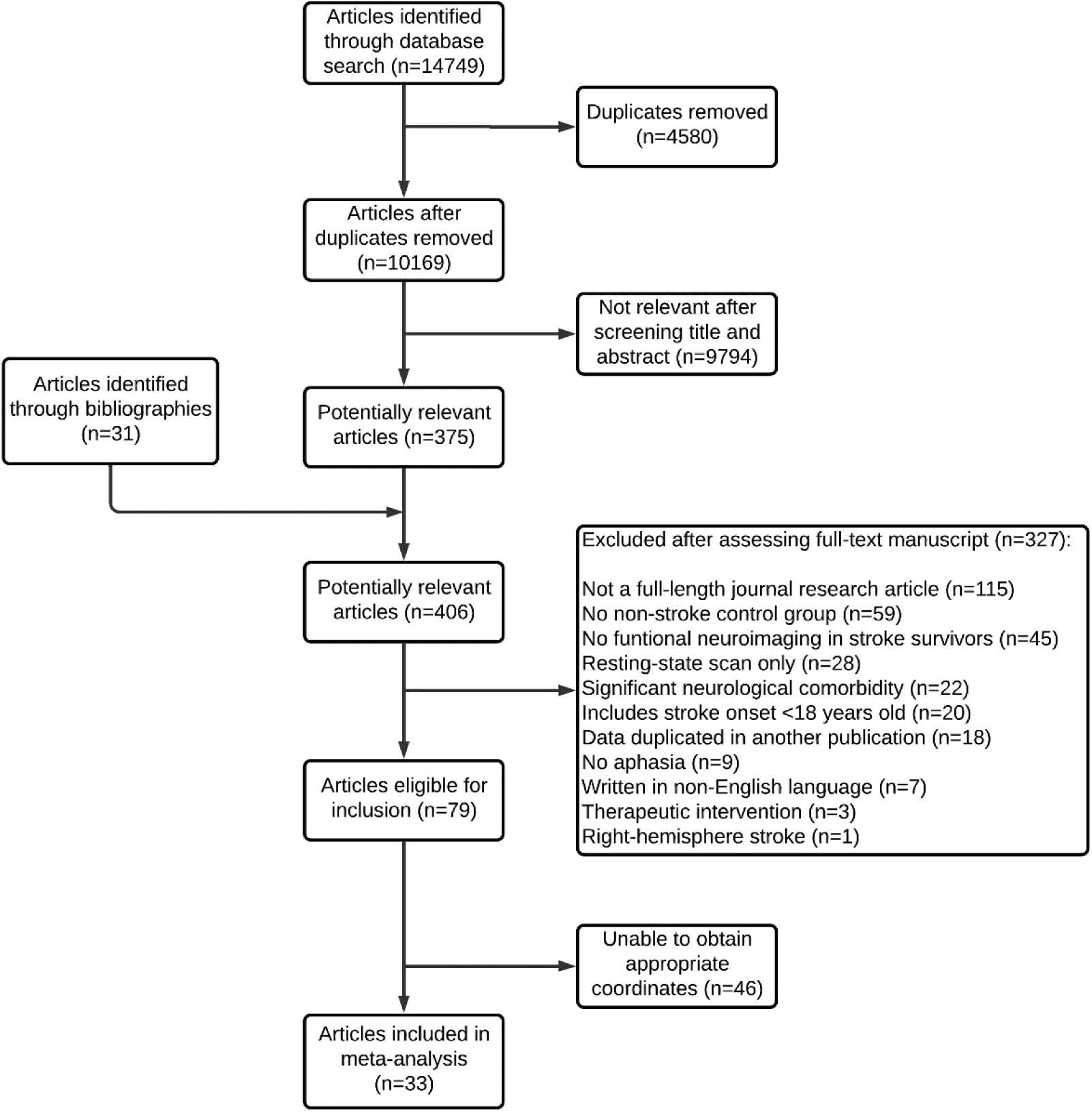
Flowchart of the selection process for included papers. Flowchart showing the selection process at each stage of the systematic search up to April 2020. Ultimately, activation foci from 33 papers were included in the ALE analysis.

The 64 PSA groups did not have significantly different mean ages compared to the 37 control groups (median 57.4 [IQR 9.0] years in PSA vs. 57.0 [IQR 8.2] years in controls; Mann-Whitney U-test, U=878, two-sided p=0.18). Every pair of datasets contrasted in this paper had mean ages that were not statistically significantly different (Supplementary Table S31). Fig.2 contains histograms of the mean ages of the groups.

**Figure 2:**
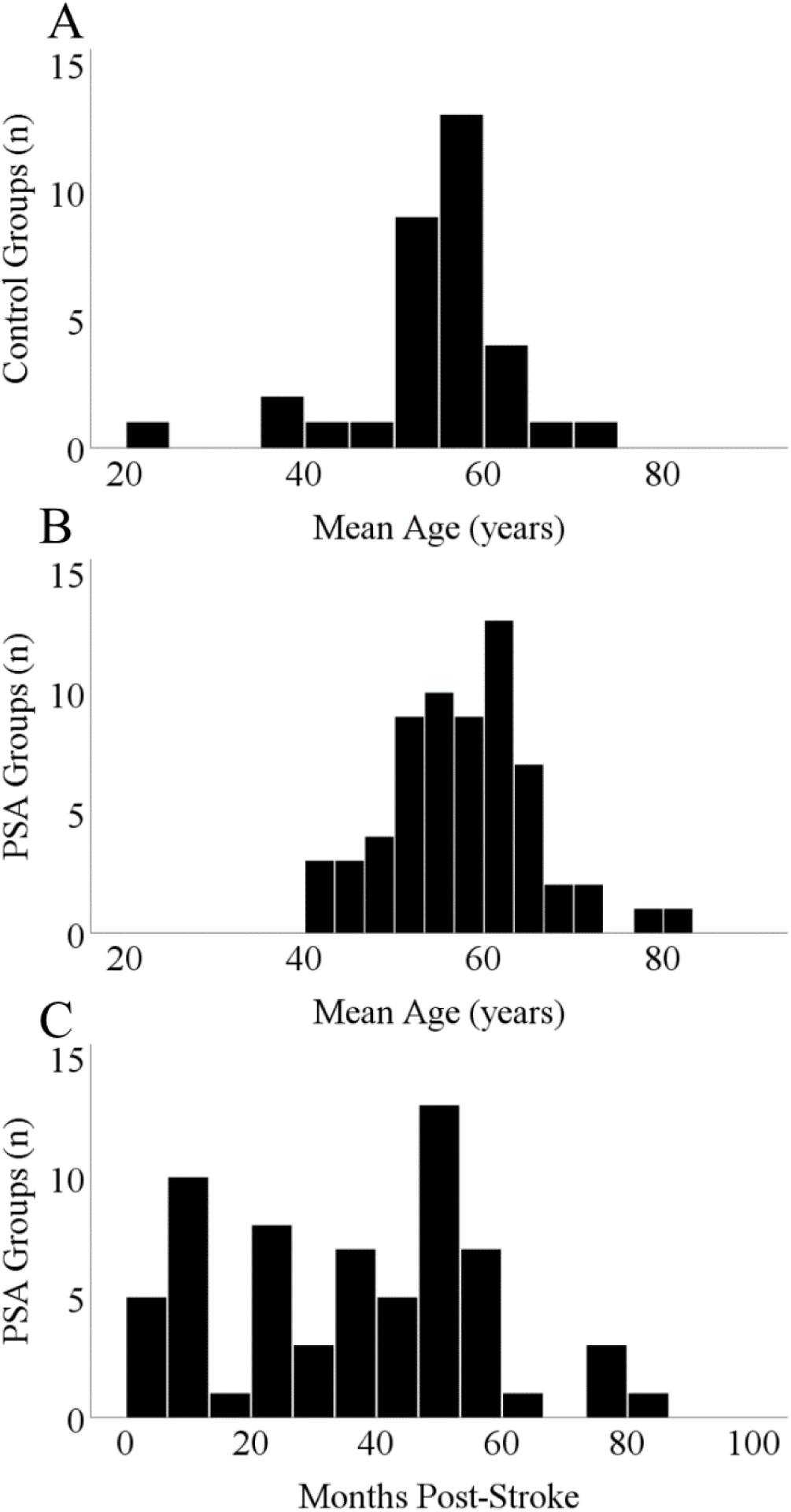
Histogram showing the distribution of participant groups with age and time post-stroke. A: Histogram showing the number of control groups for each ‘mean age’. B: Histogram showing the number of PSA groups for each ‘mean age’. C: Histogram showing the number of PSA groups for each post-stroke time period.

### Omnibus ALE analysis

The omnibus analysis compared all language tasks in all PSA against control groups (Fig.3). PSA consistently activated bilateral regions, including: left frontal lobe (inferior frontal gyrus (IFG) pars opercularis/triangularis, frontal orbital cortex, middle frontal gyrus (MFG)); left temporal lobe (posterior middle temporal gyrus (MTG)); midline cortex (superior frontal gyrus (SFG), supplementary motor cortex (SMC), paracingulate gyrus); right frontal lobe (IFG pars opercularis/triangularis, frontal orbital cortex, precentral gyrus); right insula; and right temporal lobe (posterior superior temporal gyrus (STG), Heschl’s gyrus, planum temporale) (Supplementary Table S5). A conjunction demonstrated that both PSA and control groups consistently activated overlapping regions in: left frontal lobe (frontal operculum cortex, IFG pars opercularis/triangularis, frontal orbital cortex, MFG); left temporal lobe (posterior MTG); midline cortex (SFG, SMC, paracingulate gyrus); right frontal lobe (frontal operculum, frontal orbital cortex); right temporal lobe (posterior STG); and right parietal lobe (posterior supramarginal gyrus) (Supplementary Table S7). This highlights that the language network is bilateral in both controls and PSA. Furthermore, multiple regions throughout both hemispheres were consistently activated in PSA but were also involved in language pre-morbidly rather than being recruited ‘de novo’ post-stroke.

**Figure 3:**
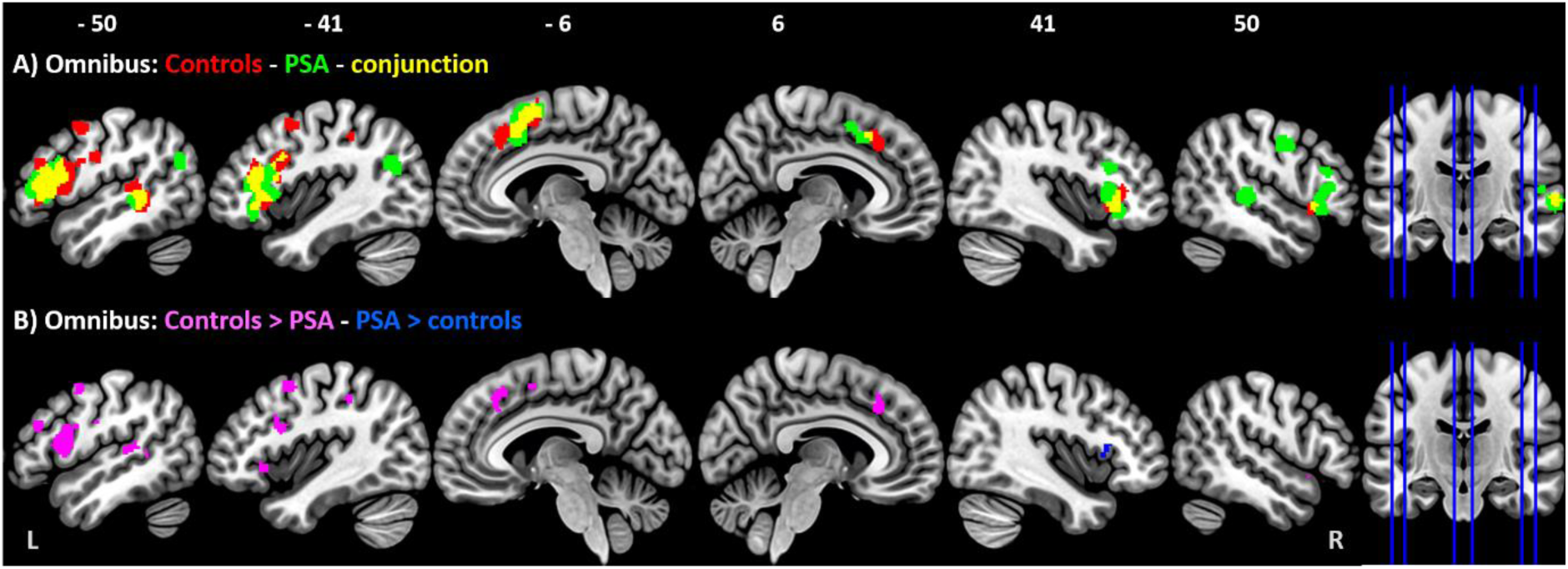
Omnibus ALE analysis for all language tasks in PSA and healthy controls. A: ALE maps of all tasks in PSA (green clusters), in controls (red clusters) and conjunction map of all tasks in both PSA and controls (yellow clusters). ALE single dataset analyses thresholded at p<0.001 uncorrected voxel-wise, FWE p<0.05 cluster wise, 1000 permutations. B: ALE maps of ‘Omnibus: controls > PSA’ (violet clusters) and ‘Omnibus: PSA > controls’ (blue clusters). ALE contrast analyses thresholded at p<0.05, 10000 permutations, minimum cluster extent 200ml.

The PSA group activated the right frontal operculum and IFG pars opercularis more consistently during language than controls (Supplementary Table S8). This cluster overlaps with the Multiple Demand (MD) network (Fedorenko *et al*., 2013), suggesting it is domain-general (Fig.7A).

Controls activated multiple regions more consistently than the PSA group, including midline SFG, SMC, and paracingulate gyrus as well as right IFG pars triangularis and right temporal pole (Supplementary Table S8). The midline SFG and paracingulate gyrus cluster overlaps with the MD network (Fedorenko *et al*., 2013), suggesting it is domain-general (Fig.7A). Since all strokes were restricted to the left hemisphere, this result demonstrates that a set of undamaged language and domain-general regions are activated less consistently in PSA than controls.

### Comprehension versus production

During comprehension tasks, PSA consistently activated regions in: left frontal lobe (IFG pars opercularis/triangularis, frontal orbital cortex); left temporal lobe (posterior MTG); midline cortex (SFG, SMC); right frontal lobe (IFG pars triangularis, frontal orbital cortex, MFG); and right insula (Fig.4A, Supplementary Table S9). Both PSA and controls consistently activated overlapping regions during comprehension in left frontal lobe (Fig.4A, IFG pars opercularis/triangularis, frontal orbital cortex) and left posterior MTG (Supplementary Table S11). PSA activated the right anterior insula more consistently during comprehension than controls (Fig.4B, Supplementary Table S12); this cluster overlaps with the MD network (Fedorenko *et al*., 2013), suggesting it is domain-general (Fig.7B). Controls activated multiple regions more consistently during comprehension than PSA, including midline cortical regions (SFG, paracingulate gyrus) that are unlikely to be lesioned following a middle cerebral artery (MCA) stroke (Fig.4B, Supplementary Table S12).

**Figure 4:**
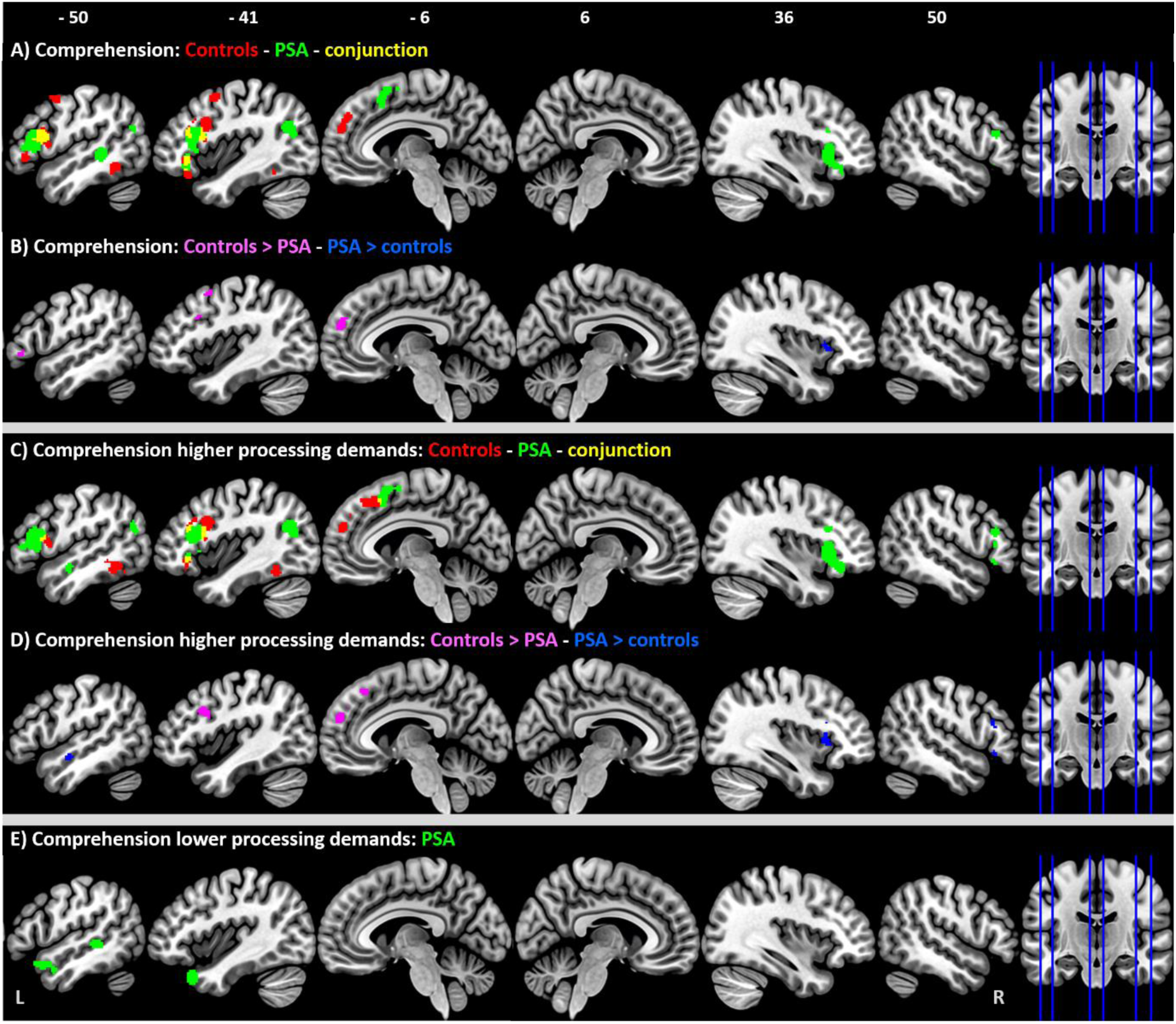
ALE analysis for comprehension tasks in PSA and healthy controls. A: ALE maps of all comprehension tasks in PSA (green clusters) and in controls (red clusters), and the conjunction map (yellow clusters). B: ALE maps of ‘Comprehension: controls > PSA’ (violet clusters) and ‘Comprehension: PSA > controls’ (blue clusters). C: ALE maps of higher processing demands comprehension tasks in PSA (green clusters), in controls (red clusters) and the conjunction map (yellow clusters). D: ALE maps of ‘High demand comprehension tasks: controls > PSA’ (violet clusters) and ‘High demand comprehension tasks: PSA > controls’ (blue clusters). E: ALE map of lower processing demand comprehension tasks in PSA only (green clusters). Panels A, C, E: ALE single dataset analyses thresholded at p<0.001 uncorrected voxel-wise, FWE p<0.05 cluster wise, 1000 permutations. Panels B, D: ALE contrast analyses thresholded at p<0.05, 10000 permutations, minimum cluster extent 200ml.

During production tasks, PSA consistently activated regions in: left frontal lobe (IFG pars triangularis); midline cortex (SFG, SMC); right frontal lobe (IFG pars triangularis, precentral gyrus); right insula; and right temporal lobe (posterior STG, Heschl’s gyrus) (Fig.5A, Supplementary Table S13). Both PSA and controls consistently activated overlapping regions during production in: left frontal lobe (IFG pars triangularis); midline cortex (SFG, SMC, paracingulate gyrus); and right temporal lobe (posterior STG) (Fig.5A, Supplementary Table S15). This highlights that multiple regions throughout both hemispheres are consistently activated during language production in PSA that were involved in language pre-morbidly rather than being recruited ‘de novo’ post-stroke. The PSA group did not activate any regions more consistently during production than controls (Supplementary Table S16). Controls activated multiple regions more consistently during production than PSA, including: midline cortex (SFG, SMC, paracingulate gyrus); right frontal lobe (frontal orbital cortex, precentral gyrus); right insula; and right temporal lobe (Heschl’s gyrus, posterior STG, temporal pole) (Fig.5B, Supplementary Table S16). Again these regions fall outside of the left MCA territory and thus were unlikely to have been lesioned by the stroke. The midline SFG and paracingulate gyrus cluster overlaps with the MD network (Fedorenko *et al*., 2013), suggesting it is domain-general (Fig.7C).

**Figure 5:**
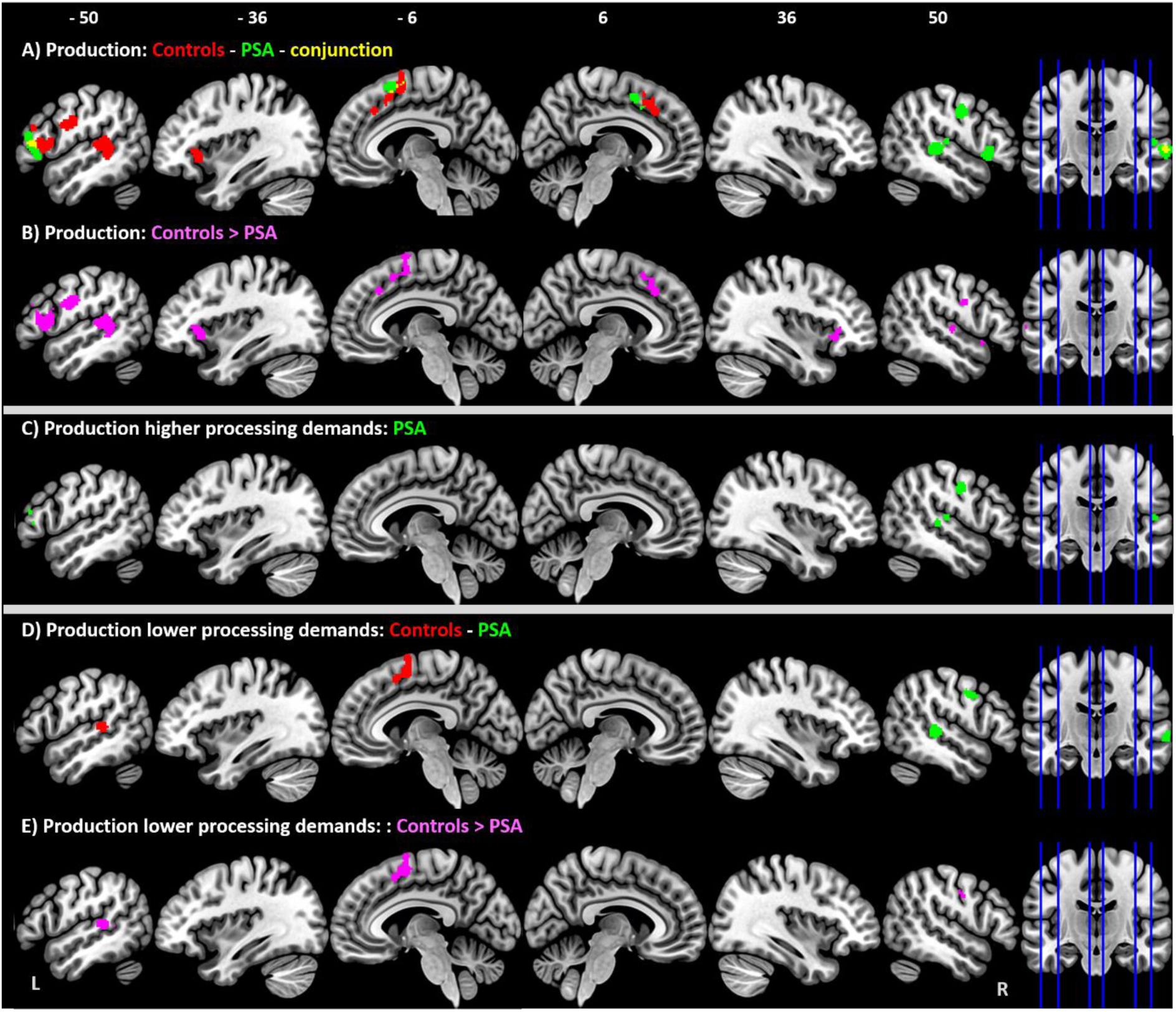
ALE analysis for production tasks in PSA and healthy controls. A: ALE maps of all production tasks in PSA (green clusters), in controls (red clusters) and conjunction map of all production tasks in both PSA and controls (yellow clusters). B: ALE maps of ‘Production tasks: controls > PSA’ (violet clusters). C: ALE map of higher processing demand production tasks in PSA (green clusters). D: ALE map of lower processing demand production tasks in PSA (green clusters) and in controls (red clusters). E: ALE map of ‘Lower processing demand production tasks: controls > PSA’ (violet clusters). Panels A, C, E: ALE single dataset analyses thresholded at p<0.001 uncorrected voxel-wise, FWE p<0.05 cluster wise, 1000 permutations. Panels B, D: ALE contrast analyses thresholded at p<0.05, 10000 permutations, minimum cluster extent 200ml.

The PSA group activated multiple regions more consistently during comprehension than production, including: left frontal lobe (IFG pars opercularis, frontal orbital cortex); left temporal lobe (posterior MTG); left parietal lobe (angular gyrus); right insula; and right frontal lobe (MFG) (Fig.6B, Supplementary Table S17). Conversely, regions of the right frontal lobe (IFG pars opercularis, precentral gyrus) and right temporal lobe (posterior STG, planum temporale) were more consistently activated during production than comprehension (Fig.6B, Supplementary Table S17). Controls activated the left frontal lobe (frontal orbital cortex, frontal pole), left temporal lobe (temporal pole, temporooccipital inferior temporal gyrus) and midline SFG/paracingulate gyrus more consistently during comprehension than production (Fig.6A, Supplementary Table S18). Conversely, controls activated the left frontal lobe (IFG pars opercularis/triangularis, frontal orbital cortex, precentral gyrus), left insula, left temporal lobe (planum temporale, temporooccipital MTG), left parietal lobe (posterior supramarginal gyrus), midline cortex (SFG, SMC, paracingulate gyrus) and right temporal lobe (temporal pole, posterior STG) more consistently during production than comprehension (Fig.6A, Supplementary Table S18).

**Figure 6:**
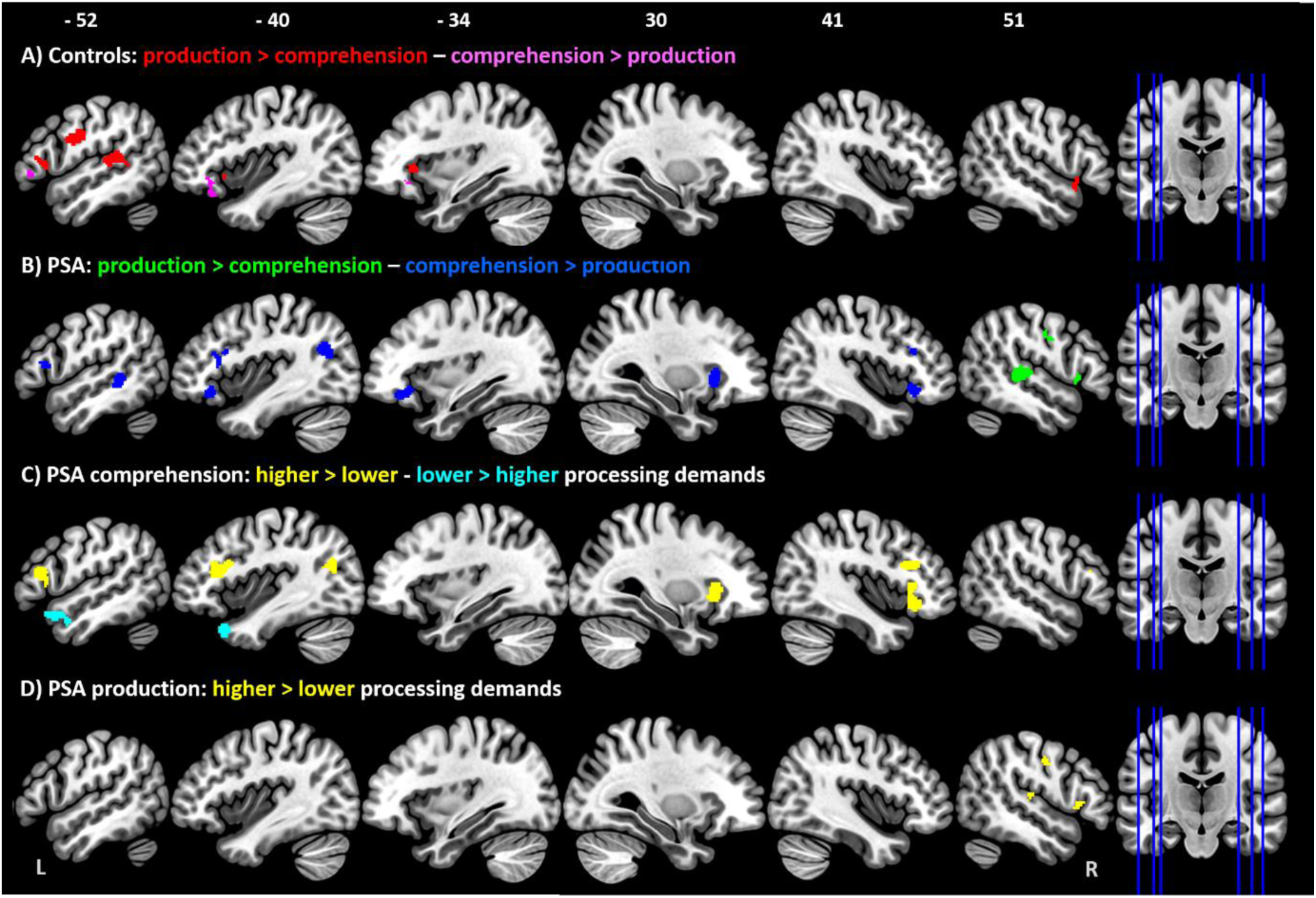
ALE analysis of within group task differences. A: ALE maps of ‘controls: production > comprehension tasks’ (red clusters), and ‘controls: comprehension > production tasks’ (violet clusters). B: ALE maps of ‘PSA: production > comprehension tasks’ (green clusters), and ‘PSA: comprehension > production tasks’ (blue clusters). C: ALE maps of ‘PSA comprehension tasks: higher > lower processing demands’ (yellow clusters) and ‘PSA comprehension tasks: lower > higher processing demands’ (cyan clusters). D: ALE maps of ‘PSA production tasks: higher > lower processing demands’ (yellow clusters). ALE contrast analyses thresholded at p<0.05, 10000 permutations, minimum cluster extent 200ml.

### Higher versus lower processing demand

Both PSA and control groups consistently activated overlapping regions in left frontal lobe (IFG pars opercularis, frontal orbital cortex, MFG) and midline SFG during higher demand comprehension tasks (Fig.4C, Supplementary Table S22). The PSA group activated the left anterior STG, right frontal lobe (IFG pars opercularis/triangularis, frontal orbital cortex) and right anterior insula more consistently during higher demand comprehension tasks than controls (Fig.4D, Supplementary Table S23). The cluster of more consistent activation in the left anterior STG demonstrates that aphasia recovery might involve more consistent utilisation of undamaged left hemisphere language regions post-stroke. The right anterior insular cluster overlaps with the MD network (Fedorenko *et al*., 2013), suggesting it is domain-general, while the right IFG pars opercularis/triangularis cluster overlaps with the semantic control network known to be involved during executively demanding semantic cognition in healthy individuals (Noonan *et al*., 2013). Controls activated multiple regions more consistently during higher demand comprehension tasks than in PSA, including midline cortex (SFG, paracingulate gyrus) (Fig.4D, Supplementary Table S23).

PSA activated the left frontal lobe (IFG pars opercularis/triangularis), right frontal lobe (frontal operculum cortex, IFG pars triangularis) and right anterior insula more consistently during higher demand than lower demand comprehension tasks (Fig.6C, Supplementary Table S24). These three clusters all overlap with the semantic control network known to be involved during executively demanding semantic cognition in healthy individuals (Noonan *et al*., 2013), while the right anterior insular cluster additionally overlaps with the MD network (Fedorenko *et al*., 2013), suggesting it is domain-general (Fig.7D).

**Figure 7:**
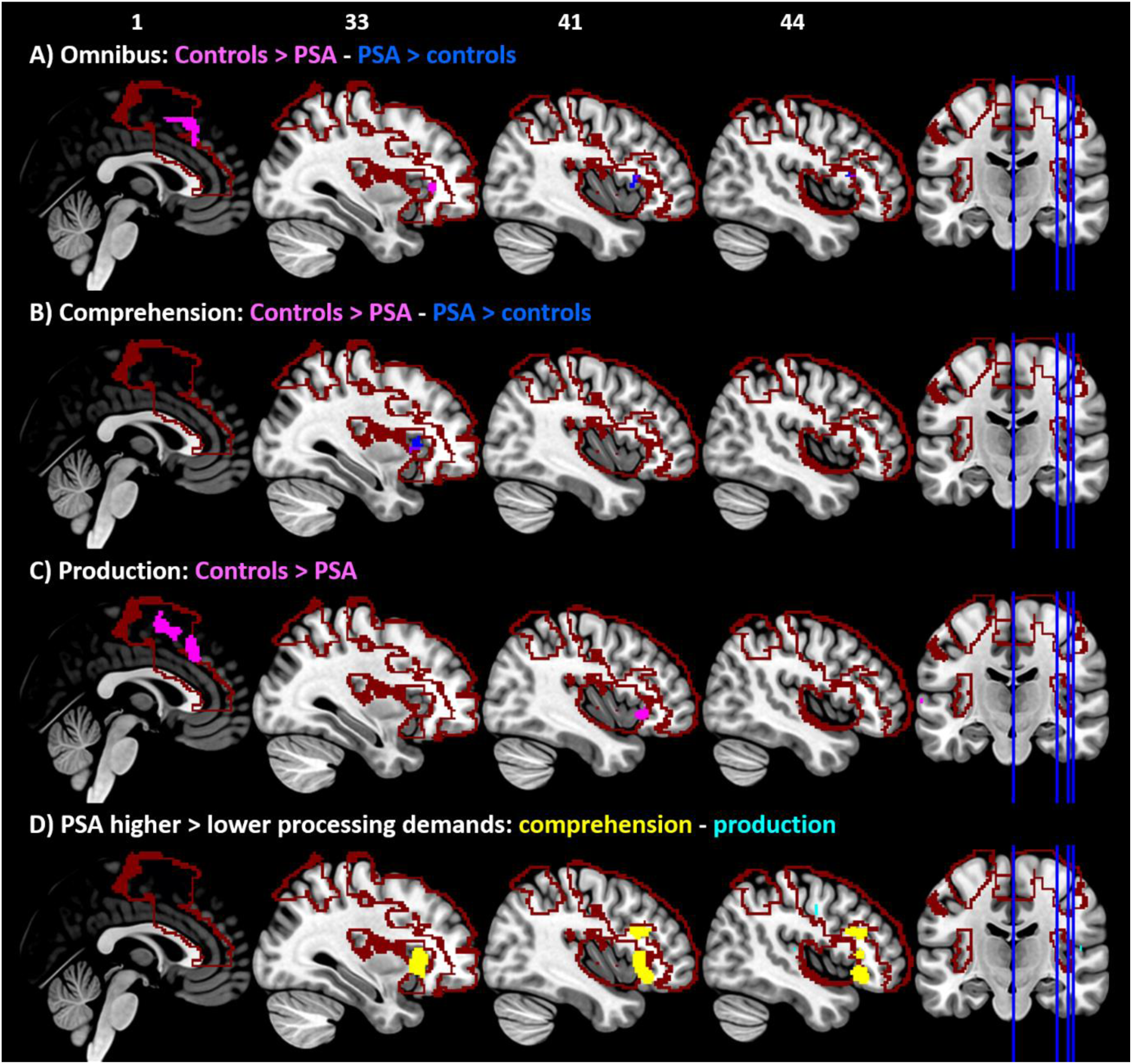
Overlaps between clusters identified in the ALE analysis and the Multiple Demand Network. A: ALE maps of ‘Omnibus analysis: controls > PSA’ (violet clusters) and ‘Omnibus analysis: PSA > controls’ (blue clusters). B: ALE maps of ‘Comprehension: controls > PSA’ (violet clusters) and ‘Comprehension: PSA > controls’ (blue clusters). C: ALE maps of ‘Production: controls > PSA’ (violet clusters). D: ALE maps of ‘Comprehension in PSA: higher> lower processing demands’ (yellow cluster) and ‘Production in PSA: higher > lower processing demands’ (cyan cluster). All panels include the outline of the Multiple Demand network (maroon) (Fedorenko *et al*., 2013). ALE contrast analyses thresholded at p<0.05, 10000 permutations, minimum cluster extent 200ml.

Only 110 foci were obtained from 78 controls in 7 participant groups performing lower demand comprehension tasks. Accordingly, there were too few groups to perform ALE meta-analyses contrasting higher versus lower demand comprehension tasks in controls, or contrasting lower demand comprehension tasks in PSA vs. controls (Eickhoff *et al*., 2016).

PSA and controls did not consistently activate any overlapping regions during lower demand production tasks (Fig.5D, Supplementary Table S28), nor did PSA activate any regions more consistently during lower demand production tasks than controls (Supplementary Table S29). Controls activated multiple regions more consistently during lower demand production tasks than PSA, including midline SMC, right precentral gyrus and right posterior STG (Fig.5E, Supplementary Table S29). The midline SMC cluster overlaps with the MD network (Fedorenko *et al*., 2013), suggesting it is domain-general.

PSA activated the right frontal lobe (frontal operculum cortex, IFG pars opercularis, precentral gyrus) and right temporal lobe (planum temporale, Heschl’s gyrus) more consistently during higher demand than lower demand production tasks (Fig.6D, Supplementary Table S30). The right precentral gyrus cluster and right IFG pars opercularis/frontal operculum cluster both overlapped with the MD network (Fedorenko *et al*., 2013), suggesting they are domain-general.

Only 189 foci were obtained from 185 controls in 8 groups performing higher demand production tasks. Accordingly, there were too few groups to perform ALE meta-analyses contrasting higher versus lower demand production tasks in controls, or contrasting higher demand production tasks in PSA vs. controls (Eickhoff *et al*., 2016).

### Time post-stroke

The literature is strongly biased as most PSA underwent neuroimaging in the chronic phase post-stroke. The 64 PSA groups had median times post-stroke of 38.0 (IQR 34.5) months (Fig. 2). Only five papers, representing six of the 64 PSA groups, repeated functional neuroimaging longitudinally at multiple timepoints (Cardebat *et al*., 2003; Radman *et al*., 2016; Long *et al*., 2017; Nenert *et al*., 2018; Stockert *et al*., 2020). When counting the ‘earliest’ timepoint at which each PSA group was scanned, only 9/64 groups had mean times post-stroke less than 6 months (Cardebat *et al*., 2003; Mattioli *et al*., 2014; Geranmayeh *et al*., 2016; Radman *et al*., 2016; Long *et al*., 2017; Qiu *et al*., 2017; Nenert *et al*., 2018; Stockert *et al*., 2020). Accordingly, there were too few groups to contrast PSA before versus after six months (Eickhoff *et al*., 2016).

## Discussion

In order to identify the specific regions that are activated more consistently in PSA than healthy individuals, and to investigate whether there are consistent differences in activation across different language tasks and between recovery timepoints, we performed a large-scale ALE analysis of functional neuroimaging studies in PSA. We obtained coordinate-based functional neuroimaging data for 481 PSA, which is over four times larger than the last ALE analysis on this topic (n=105) (Turkeltaub *et al*., 2011). The results provide novel insights into the mechanisms underlying language network changes post-stroke that might hitherto have been obscured by the limited sample size of any individual study in this area.

### Main findings

PSA activated various regions of the right anterior insula and IFG more consistently than controls: across all language tasks (frontal operculum, IFG pars opercularis); during comprehension tasks (anterior insula); and during higher demand comprehension tasks (IFG pars opercularis/triangularis, frontal orbital cortex and anterior insula). These right anterior insular/IFG regions seem to be implicated in task difficulty as they are activated more consistently during higher than lower demand comprehension tasks (frontal operculum, IFG pars triangularis, anterior insula) and during higher than lower demand production tasks (operculum, IFG pars opercularis). PSA activated several left hemisphere regions (IFG pars opercularis, frontal orbital cortex, posterior MTG, angular gyrus) and a smaller number of right hemisphere regions (insula and MFG) more consistently during comprehension than production tasks. Finally, PSA activated only right frontal (IFG pars opercularis, precentral gyrus) and temporal (planum temporale, posterior STG) regions more consistently during production than comprehension tasks, in keeping with a role for right superior temporal cortex during auditory feedback monitoring of produced speech (Wolpert *et al*., 1995; Rauschecker and Scott, 2009; Houde and Nagarajan, 2011; Yamamoto *et al*., 2019).

### Novel implications

#### a) Similarities between the language network in PSA and healthy individuals

A previous ALE analysis in PSA concluded that the language network in controls is left-lateralised, whereas PSA consistently activate additional right hemisphere regions that are not consistently activated in controls (Turkeltaub *et al*., 2011). The clear picture that emerges from the current, much larger ALE analysis is different in a fundamental way. Whilst one can find reliably different levels of activation likelihood between the PSA and control groups, these differences all fall within regions that are found to activate in both groups; in classical neuropsychological terminology (Shallice, 1988), there is not a classical dissociation between PSA and control groups. Thus in the omnibus language ALE analysis, the conjunction demonstrated that both PSA and controls consistently activated overlapping regions across the left and right frontal and temporal lobes, right parietal lobe, and midline cortex. Two important implications are that (a) right as well as left hemisphere areas make importance contributions to language and (b) that regions, consistently activated by language tasks in PSA, are also involved in language pre-morbidly. This runs counter to the view that these areas are recruited ‘de novo’ post-stroke.

#### b) Undamaged brain regions less consistently activated in PSA than controls

Irrespective of how the language tasks were divided (all language tasks, comprehension, production, higher demand vs. lower demand tasks), we found that in PSA certain regions are less consistently activated than in controls. These areas were not only left hemisphere regions that might have been lesioned directly by the stroke (i.e., within the left hemisphere MCA: cf. (Phan *et al*., 2005; Zhao *et al*., 2020)) but also domain-general regions of midline superior frontal and paracingulate cortex, right insular cortex and right fronto-temporal cortex. This result implies that the language and cognitive deficits observed in PSA might not be a simple reflection of the lesioned areas but might result from combinations of lesioned and under-engaged areas. Accordingly, the use of task-based fMRI may be an important addition for future studies that aim to explore the neural bases of aphasia or build prediction models (Saur *et al*., 2010; Skipper-Kallal *et al*., 2017a; van Oers *et al*., 2018). Less consistent activation in regions distant to the lesions might reflect functional diaschisis, i.e., reduced task-related engagement throughout a connected network where one or more nodes have been compromised by damage (Carrera and Tononi, 2014). Alternatively from a more functional viewpoint, these distant regions may be less engaged because in PSA language is performed sub-optimally and therefore the full extent of the distributed language network is under-utilised.

#### c) Implications for mechanisms of post-stroke aphasia recovery

*Neurocomputational invasion* would predict that the post-stroke language network should expand to include novel non-language regions that were not consistently activated in healthy individuals (Keidel *et al*., 2010; Stefaniak *et al*., 2020). This mechanism is complementary to the classical notion that right hemisphere homologues of left hemisphere language regions are quiescent in health but become activated to perform similar language computations following left hemisphere stroke (Finger *et al*., 2003; Turkeltaub *et al*., 2011). A second linked idea is the notion of transcallosal disinhibition (Marshall, 1984; Heiss and Thiel, 2006). This proposes that right hemisphere, homologous regions are quiescent in health because they are inhibited transcallosally by the dominant left hemisphere, but can be ‘released’ when these dominant areas are damaged. This idea has been an important motivation for trials of non-invasive brain stimulation to inhibit the right IFG pars triangularis to aid language recovery through a shift back to left hemisphere areas (Ren *et al*., 2014; Bucur and Papagno, 2019). Previous work (Stefaniak *et al*., 2020) has noted that these hypotheses appear to be biologically-expensive (areas are maintained but not used, except in people who happen to have the right type and location of damage), computationally underspecified (e.g., how right hemisphere regions can develop language functions when they are being constantly inhibited), and are an untested extension of findings from low-level, non-language motor circuitry (Ferbert *et al*., 1992; Di Lazzaro *et al*., 1999). Additional counter evidence includes: chronic language weaknesses can be found following right hemisphere damage (Gajardo-Vidal *et al*., 2018); and, residual language abilities in PSA have been related to the level of right hemisphere activation (Crinion and Price, 2005; Griffis *et al*., 2017; Skipper-Kallal *et al*., 2017b). The current study adds to these observations in that multiple regions throughout both hemispheres are consistently activated during language in both PSA and controls. Looking across these studies, it would seem that there is a solid empirical basis to move beyond oversimplified discussions of ‘left versus right’ language lateralisation and, instead, to explore how a bilateral, albeit asymmetrically left-biased, language network supports healthy function and generates aphasia after damage and partial recovery.

*Variable neurodisplacement* postulates that aphasia recovery involves increased utilisation of spare capacity within regions that are part of the premorbid language network but downregulated in health to save neural resources. Dynamic responses to performance demands in health and after damage could involve upregulation of language-specific and/or domain-general executive functions (Stefaniak *et al*., 2020). Accordingly, variable neurodisplacement encompasses the hypothesis that increased utilisation of domain-general executive regions aids language recovery post-stroke (Sharp *et al*., 2010; Geranmayeh *et al*., 2014). As noted above, a key finding from these ALE analyses was that many, bilateral regions were commonly engaged by PSA and control groups. Even where there were graded differences in favour of PSA over controls (e.g., more consistent activation in the right anterior insula and IFG), these are consistent with enhanced utilisation of demand-control regions due to increased task difficulty rather than ‘expansion’ into new territory via neurocomputational invasion. Thus, in the PSA group, there was more consistent activation of the right anterior insula/operculum and IFG during higher than lower demand comprehension and production tasks. Although there was insufficient control data for performance demand to be examined in this ALE analysis, these same right anterior insula/IFG regions are known to be recruited during difficult tasks in healthy individuals: the right IFG has been implicated in domain-general top-down control in health (Koechlin and Jubault, 2006; Meinzer *et al*., 2012; Baumgaertner *et al*., 2013); a previous ALE meta-analysis found that effortful listening under difficult conditions in healthy individuals is associated with consistent activation in the bilateral insulae (Alain *et al*., 2018); and all ALE-identified right hemisphere regions overlap with either domain-general regions of the MD network (Fedorenko *et al*., 2013) or regions of the semantic control network known to be involved during executively-demanding semantic cognition in healthy individuals (Noonan *et al*., 2013).

The results do not suggest that there is a global, undifferentiated upregulation of all domain-general neural resources in PSA. Indeed, we repeatedly found less consistent activation in midline regions of the SFG/paracingulate gyrus in PSA compared to controls, including during higher demand comprehension tasks. These midline clusters overlap with at least some definitions of the domain-general executive network (Fedorenko *et al*., 2013). In contrast to our findings, increased activation in the same midline region has been associated with language recovery between two weeks and four months post-stroke (Geranmayeh *et al*., 2017). It is not clear what the basis of these opposing results is, but one possibility is that this ALE analysis was predominantly based on data collected from patients in the very chronic (see below) rather than sub-acute stage. If correct, it may be the case that the executive functions supported by medial prefrontal regions (e.g., response conflict, task planning (Dosenbach *et al*., 2008; Mansouri *et al*., 2017)) are critical during early phases of recovery when performance is at its most impaired, but in relatively well-recovered, chronic PSA these mechanisms are not required (indeed continued involvement might signal poor recovery).

#### d) Age of PSA included in functional neuroimaging studies

As is commonly the case in stroke research (Fareed *et al*., 2012; Thomalla *et al*., 2017), the median ages of the 64 included PSA subject groups was lower (57.4 years) than the average stroke patient (e.g., the median age of the UK stroke population was 77 in 2017 (SSNAP, 2017). This may limit the generalisability of results obtained from functional neuroimaging studies to the ‘real-world’, and future studies should investigate patterns of activation in older PSA that are more representative of the average stroke survivor.

### Unanswered questions: time-dependent changes and longitudinal imaging studies

We identified areas of enquiry that have had little attention in the literature to date. It was not possible to ascertain whether there are consistent activation differences between subacute and chronic PSA. The 64 PSA groups had median times post-stroke of 38.0 months and even when counting the ‘earliest’ timepoint at which each PSA group was scanned, only 9/64 PSA groups were less than 6 months post-stroke. This dearth of data meant it was not possible to use ALE to explore differences between sub-acute and chronic PSA. Importantly, this indicates a pressing need for future studies of this early period, when there is the fastest rate of language recovery (Pedersen *et al*., 1995; Yagata *et al*., 2017). Additionally, it was not possible to explore longitudinal fMRI changes given the extremely limited number of longitudinal PSA fMRI studies. Even among papers that reported longitudinal information, several were small (n<10 participants) and there was considerable variation with respect to which language or non-language cognition was explored and the timing of the first imaging timepoint (from the first few days to a few months post-stroke). The relative lack of studies and small sample sizes are unsurprising given the considerable logistic challenges involved in imaging subacute stroke patients. However, longitudinal studies are a powerful approach for exploring the neural bases of recovery (because the different starting points and inter-participant variations are controlled), and particularly for exploring whether language network changes observed in the chronic phase occur immediately or over time. Such information will be critical for understanding the mechanisms underpinning both instantaneous resilience to the effects of damage, degeneracy, and longer-term experience-dependent plasticity (Price and Friston, 2002; Ueno *et al*., 2011; Chang and Lambon Ralph, 2020; Sajid *et al*., 2020; Stefaniak *et al*., 2020).

## Conclusion and limitations

The results of this large-scale analysis argue against classical neurocomputational invasion accounts of PSA language, i.e., expansion of the language network post-damage into new territories. Instead, (a) there is considerable overlap between the bilateral language-related functional networks observed in PSA and controls; (b) the PSA participants are less likely than controls to activate certain regions including areas beyond their core lesions in the left MCA territory; and (c) are more likely to engage executive-control related regions of the right anterior insula and IFG. These results fit with a view that language is supported by a dynamic, bilateral albeit left-asymmetric network, and consistent with the variable neurodisplacement hypothesis. The size of this (random-effects) analysis (including data pertaining to 481 PSA with a heterogenous variety of lesion locations and aphasia profiles), should mean that the results will generalise to the wider patient population.

Despite its size and clear results, inevitably this study has limitations. First, we did not have information regarding lesion location and thus were unable to investigate how this might influence activation patterns post-stroke (Stockert *et al*., 2020). Second, it is important to be cautious about decreased neurovascular coupling post-stroke, which itself could generate false differences between patients and controls. However, altered neurovascular coupling is less likely in chronic patients and in undamaged regions of the midline and right hemisphere (Geranmayeh *et al*., 2015). Third, as is typical, the current analysis is based on mass-univariate BOLD activation differences. It is possible that ‘neural reprogramming’ post-stroke might entail differences in utilisation that are only observable using connectivity analyses (Schofield *et al*., 2012; Meier *et al*., 2018) or multivariate techniques (Fischer-Baum *et al*., 2017; Lee *et al*., 2017). Currently, there are very few studies that have used such techniques in PSA. Finally, domain-general upregulation might occur at earlier stages post stroke, as has been suggested by the largest longitudinal study in PSA to date (Geranmayeh *et al*., 2017). However, this analysis demonstrates that there are currently insufficient acute and sub-acute data in the published literature to investigate time-dependent changes at earlier stages post-stroke.

### Acknowledgements

We would like to thank Sonia Brownsett, Fatemeh Geranmayeh, Glyn Hallam, Jessica Hodgson, Narges Radman, Michael Mouthon, Alex Leff, Stephen Wilson, Yuan Tao, Erin Meier, and Elena Barbieri for kindly sharing unpublished coordinates relating to their published studies for use in this ALE meta-analysis.

