## Supplementary Material for "Language networks in aphasia and health: a 1000 participant Activation Likelihood Estimate analysis"

Supplementary Methods:

Full search strategy and inclusion criteria

The databases Medline, Embase and PsycINFO were searched (using OvidSP for Medline and Embase) with the following criteria:

(aphasia OR dysphasia OR language OR fluency OR phonology OR semantics OR naming OR repetition OR comprehension OR speaking) AND (stroke OR ischaemia OR ischemia OR infarct) AND (fMRI OR PET OR neuroimaging OR imaging OR functional)

A search was performed with these criteria on 13/09/2017; we considered articles published at any time before this search date. This was updated on 17/03/2018 and again on 13/04/2020. After removal of duplicates, the title and abstracts of all search results were screened for potential relevance. Full-length manuscripts of all potentially relevant articles were reviewed. In addition, the bibliographies of all potentially relevant manuscripts were reviewed for additional articles that might have been missed in the primary search.

We identified eligible articles reporting observational studies that had: a) more than one person with language impairment at any time following left hemispheric stroke; b) more than one healthy control; and c) performed BOLD fMRI or ^15^O-PET during a language task-based functional neuroimaging paradigm. Stroke survivors of both sexes, of any ethnicity, based in any clinical setting, with history of language impairment after haemorrhagic or ischaemic stroke were included. Studies with stroke survivors of any age were included as long as all aphasic group coordinates pertained to individuals who had stroke onsets when they were adults (>18 years old). Articles in which post-stroke aphasia was not specifically mentioned but in which stroke survivors had a documented language deficit at any time post-stroke were included. Studies in which stroke survivors initially had language impairments but whose language abilities had completely recovered were still included. We included studies regardless of blinding status. Conference abstracts, presentations, theses, and articles not written in English were excluded. We excluded studies without a comparator group or that related solely to alexia, agraphia, amusia, dysarthria, dysphonia, apraxia of speech or non-linguistic auditory processing but not aphasia. Articles using separately published functional neuroimaging data from controls were included as long as the control participant demographics and neuroimaging results were obtainable and the functional neuroimaging data from the aphasic group was novel. If the same stroke survivors with aphasia were included in functional neuroimaging tasks in multiple papers, only the paper containing the largest number of participants was included. The only exceptions were: (1) if exactly the same aphasic and control participants were included in two papers and performed different functional neuroimaging tasks at the same time post-stroke that were analysed identically in the two papers, in which case the two papers were considered to be different functional neuroimaging tasks of the same group and were both included; or (2) if individual participant-level peak activation coordinates were available, in which case aphasia participants that were included in both papers were excluded from one of the papers.

We then extracted peak coordinate data from the list of articles that were eligible for inclusion in our meta-analysis. We extracted coordinate data for inclusion in this ALE meta-analysis that: related to activation (not deactivation) during a language task-based functional neuroimaging experiment; was provided in standard space; was derived from whole-brain mass-univariate analyses without region of interests (ROIs), small volume corrections (SVC), or conjunctions (Müller *et al.*, 2018); and was calculated using the same significance thresholds in the aphasic group and control group. To be included, functional neuroimaging coordinates from the ‘aphasic group’ had to: exclude survivors of a right-sided stroke; exclude people without history of language impairment; and be collected before any study-specific research intervention might have occurred. Imaging analyses could not use masking, other than exclusive lesion masking or inclusive grey matter masking. In order to be as inclusive of the published literature as possible, we included coordinates obtained at any statistical significance threshold provided the same threshold was used in the aphasic group and control group of that article. If coordinates meeting these criteria for both the aphasic group and control group were not provided in the publication, authors were emailed for unpublished coordinates.

Coordinates from tasks at different timepoints on the same participant group were not pooled; only tasks performed at the longest timepoint post-stroke for each group were included. This was because we expect language network changes to occur between subacute and chronic stages in the same group, and we predicted that most functional neuroimaging studies would have been performed in the chronic phase post-stroke after language change had slowed or plateaued.

Subgroup ALE meta-analyses

PSA groups were divided into one of two categories according to a characteristic of its PSA participants or a characteristic of the imaging task performed. A contrast ALE meta-analysis compared the resultant two categories of PSA groups. Each control group was divided according to how its corresponding PSA group had been categorised and a contrast meta-analysis performed on the resultant two categories of controls. If different PSA groups from within the same article possessed different patient characteristics and were divided into different categories, both groups were included in their respective categories and the paper’s control coordinates were included in both categories for the control contrast meta-analysis. If the same PSA group performed multiple imaging tasks which were divided into different categories, the coordinates for both imaging tasks were included in their respective categories. Since contrast analyses were designed to look for regions of significantly different convergence between groups, the inclusion of coordinates from the same group in both subgroup datasets being contrasted would, if anything, reduce the likelihood of finding differences and thus should not increase the false positive rate.

Supplementary Table S1: Number of articles included and excluded at each stage of the systematic search

| Search date | 13/09/2017 | 17/03/2018 | 13/04/2020 | Total |
| --- | --- | --- | --- | --- |
| Number of articles after search of Medline and Embase | 9221 (+9221) | 746 (+746) | 2095 (+2095) | 12062 (+12062) |
| Number of articles after search of PsycINFO | 11691 (+2470) | 955 (+209) | 2103 (+8) | 14749 (+2687) |
| Number of articles after removal of duplicates | 8015 (-3676) | 363 (-592) | 1791 (-312) | 10169 (-4580) |
| Number of potentially relevant articles after screening title and abstract | 317 (-7698) | 21 (-342) | 37 (-1754) | 375 (-9794) |
| Number of potentially relevant articles after screening bibliographies | 345 (+28) | 21 (0) | 40 (+3) | 406 (+31) |
| Number of articles eligible for inclusion after assessment of full-length manuscript | 67 (-278) | 2 (-19) | 10 (-30) | 79 (-327) |
| Number of articles included in meta-analysis (appropriate coordinates obtained) | 25 (-42) | 2 (0) | 6 (-4) | **33** (-46) |

Numbers outside of parentheses represent the running total of the number of articles included at the corresponding row’s stage of the systematic search, for the search date indicated by the corresponding column. Numbers in parentheses represent the number of articles added or removed from the systematic search between the previous and current row.

Supplementary Table S2: Reasons for exclusion of potentially relevant articles after reading full length manuscripts

| Search date | 13/09/2017 | 17/03/2018 | 13/04/2020 | Total |
| --- | --- | --- | --- | --- |
| Not a full-length journal research article | 102 | 7 | 6 | 115 |
| No non-stroke control group | 50 | 3 | 6 | 59 |
| No functional neuroimaging in stroke survivors | 35 | 3 | 7 | 45 |
| Resting-state only | 22 | 4 | 2 | 28 |
| Significant neurological comorbidity | 20 | 0 | 2 | 22 |
| Includes stroke onset <18 years old | 19 | 1 | 0 | 20 |
| Data duplicated in another publication | 14 | 1 | 3 | 18 |
| No aphasia | 8 | 0 | 1 | 9 |
| Written in non-English language | 6 | 0 | 1 | 7 |
| Therapeutic intervention | 1 | 0 | 2 | 3 |
| Right-hemisphere stroke | 1 | 0 | 0 | 1 |

Supplementary Table S3: Table of all PSA groups included in the ALE analysis

| Reference | Digital Object Identifier | Non-aphasic subgroup | Number in non-aphasic group | Non-aphasic group average age | PSA subgroup | Number in PSA group | PSA group average age | Months post stroke | Imaging task | Imaging modality | Task type | Task processing requirements |
| --- | --- | --- | --- | --- | --- | --- | --- | --- | --- | --- | --- | --- |
| Allendorfer 2012 | [10.12659/MSM.882518](https://dx.doi.org/10.12659%2FMSM.882518) | (only one non-aphasic group) | 32 | 51.1 | (only one PSA group) | 16 | 54.4^a^ | 44.4 | Overt verb generation>Noun repetition | fMRI | Production | High |
|  |  |  |  |  |  |  |  |  | Overt>Covert verb generation | fMRI | Production | High |
| Crinion 2005 | [10.1093/brain/awh659](https://doi.org/10.1093/brain/awh659) | (only one non-aphasic group) | 18 | 58 | Temporal patients | 8 | 57.8^b^ | 40.8 | Listening to meaningful stories>Meaningless reversed speech | fMRI | Comprehension | Low |
| Crinion 2005 | [10.1093/brain/awh659](https://doi.org/10.1093/brain/awh659) | (only one non-aphasic group) | (18) | (58) | Aphasic control patients | 9 | 66^b^ | 48.1 | (Listening to meaningful stories>Meaningless reversed speech) | (fMRI) | (Comprehension) | (Low) |
| Szaflarski 2011 | 10.1016/j.jstrokecerebrovasdis.2010.02.003 | (only one non-aphasic group) | 4 | 46 | (only one PSA group) | 4 | 53.3^a^ | 57 | Picture-name matching > Determining whether two geometric figures are identical | fMRI | Comprehension | Low |
| Mattioli 2014 | 10.1161/STROKEAHA.113.003192 | (only one non-aphasic group) | 10 | (unknown) | (only one PSA group) | 12 | 64.1^b^ | 0.08 | Intelligible sentence comprehension>Unintelligible sentence comprehension | fMRI | Comprehension | High |
| Qiu 2017 | 10.4103/1673-5374.198996 | (only one non-aphasic group) | 10 | 55.89 | (only one PSA group) | 10 | 55.9^a^ | 1-3 | Picture naming>Baseline | fMRI | Production | Low |
| Robson 2014 | 10.1093/brain/awt373 | (only one non-aphasic group) | 12 | 71 | (only one PSA group) | 12 | 70.1^b^ | 20.3 | Picture animate-inanimate judgement>Dual-baseline | fMRI | Comprehension | High |
|  |  |  |  |  |  |  |  |  | Word animate-inanimate judgement>Dual-baseline | fMRI | Comprehension | High |
| Skipper-Kallal 2017 | 10.1155/2017/8740353 | (only one non-aphasic group) | 37 | 58.7 | (only one PSA group) | 39 | 59.8^b^ | 52.9 | Covert naming>Fixation | fMRI | Production | Low |
|  |  |  |  |  |  |  |  |  | Overt naming>Fixation | fMRI | Production | Low |
| Blank 2003 | [10.1002/ana.10656](https://doi.org/10.1002/ana.10656) | (only one non-aphasic group) | 12 | 52.5 | POp+ | 7 | 50^a^ | 39 | Propositional speech>Listening to environmental sounds | PET | Production | High |
|  |  |  |  |  |  |  |  |  | Propositional speech>Counting aloud | PET | Production | High |
| Blank 2003 | [10.1002/ana.10656](https://doi.org/10.1002/ana.10656) | (only one non-aphasic group) | (12) | (52.5) | POp- | 7 | 61^b^ | 17 | (Propositional speech>Listening to environmental sounds) | (PET) | (Production) | (High) |
|  |  |  |  |  |  |  |  |  | (Propositional speech>Counting aloud) | (PET) | (Production) | (High) |
| Cardebat 2003 | [10.1161/01.STR.0000099965.99393.83](https://doi.org/10.1161/01.STR.0000099965.99393.83) | (only one non-aphasic group) | 6 | 50.6 | (only one PSA group) | 8 | 58.4^b^ | 1.9 (first timepoint) | Word generation>Rest | PET | Production | High |
|  |  |  |  |  |  |  |  | 11.7 (last timepoint) |  |  |  |  |
| Specht 2009 | 10.1016/j.neuroimage.2009.06.011 | (only one non-aphasic group) | 12 | (unknown) | (only one PSA group) | 12 | 49.8^a^ | 22.9 | Pseudoword decision task>Nonword decision task | PET | Comprehension | High |
| Weiller 1995 | [10.1002/ana.410370605](https://doi.org/10.1002/ana.410370605) | (only one non-aphasic group) | 6 | 35 | (only one PSA group) | 6 | 57.7^b^ | 43.2 | Verb generation>Rest | PET | Production | High |
|  |  |  |  |  |  |  |  |  | Verb generation>Pseudoword repetition | PET | Production | High |
|  |  |  |  |  |  |  |  |  | Pseudoword repetition>Rest | PET | Production | High |
|  |  |  |  |  |  |  |  |  | Pseudoword repetition>Verb generation | PET | Production | High |
| Fridriksson 2009 | 10.1002/hbm.20683 | (only one non-aphasic group) | 10 | 58.3 | (only one PSA group) | 11 | 58.8^b^ | 37.6 | Picture naming>Abstract colour picture viewing | fMRI | Production | Low |
| Warren 2009 | 10.1093/brain/awp270 | (only one non-aphasic group) | 11 | 54.6 | (only one PSA group) | 16 | 65.8^b^ | 28.8 | Listening to intelligible speech>Reversed speech | PET | Comprehension | Low |
| Sebastian 2012 | 10.1016/j.jneuroling.2012.01.003 | Normal Control 1 | 1 | 57 | Aphasic participant 1 | 1 | 60^b^ | 76 | Semantic judgement>Size judgement | fMRI | Comprehension | High |
| Sebastian 2012 | 10.1016/j.jneuroling.2012.01.003 | Normal Control 2 | 1 | (57) | Aphasic participant 2 | 1 | 53^a^ | 12 | (Semantic judgement>Size judgement) | (fMRI) | (Comprehension) | (High) |
| Sebastian 2012 | 10.1016/j.jneuroling.2012.01.003 | Normal Control 3 | 1 | (57) | Aphasic participant 3 | 1 | 53^a^ | 13 | (Semantic judgement>Size judgement) | (fMRI) | (Comprehension) | (High) |
| Sebastian 2011 | 10.1080/02687038.2011.557436 | (only one non-aphasic group) | 8 | (unknown) | Aphasic participant 1 | 1 | 62^b^ | 52 | Semantic judgement>Size judgement | fMRI | Comprehension | High |
|  |  |  |  |  |  |  |  |  | Picture naming>Scrambled picture viewing | fMRI | Production | Low |
| Sebastian 2011 | 10.1080/02687038.2011.557436 | (only one non-aphasic group) | (8) | (unknown) | Aphasic participant 2 | 1 | 57^a^ | 36 | (Semantic judgement>Size judgement) | (fMRI) | (Comprehension) | (High) |
|  |  |  |  |  |  |  |  |  | (Picture naming>Scrambled picture viewing) | (fMRI) | (Production) | (Low) |
| Sebastian 2011 | 10.1080/02687038.2011.557436 | (only one non-aphasic group) | (8) | (unknown) | Aphasic participant 3 | 1 | 60^b^ | 78 | (Semantic judgement>Size judgement) | (fMRI) | (Comprehension) | (High) |
|  |  |  |  |  |  |  |  |  | (Picture naming>Scrambled picture viewing) | (fMRI) | (Production) | (Low) |
| Sebastian 2011 | 10.1080/02687038.2011.557436 | (only one non-aphasic group) | (8) | (unknown) | Aphasic participant 4 | 1 | 40^a^ | 30 | (Semantic judgement>Size judgement) | (fMRI) | (Comprehension) | (High) |
|  |  |  |  |  |  |  |  |  | (Picture naming>Scrambled picture viewing) | (fMRI) | (Production) | (Low) |
| Sebastian 2011 | 10.1080/02687038.2011.557436 | (only one non-aphasic group) | (8) | (unknown) | Aphasic participant 5 | 1 | 70^b^ | 36 | (Semantic judgement>Size judgement) | (fMRI) | (Comprehension) | (High) |
|  |  |  |  |  |  |  |  |  | (Picture naming>Scrambled picture viewing) | (fMRI) | (Production) | (Low) |
| Sebastian 2011 | 10.1080/02687038.2011.557436 | (only one non-aphasic group) | (8) | (unknown) | Aphasic participant 6 | 1 | 51^a^ | 38 | (Semantic judgement>Size judgement) | (fMRI) | (Comprehension) | (High) |
|  |  |  |  |  |  |  |  |  | (Picture naming>Scrambled picture viewing) | (fMRI) | (Production) | (Low) |
| Sebastian 2011 | 10.1080/02687038.2011.557436 | (only one non-aphasic group) | (8) | (unknown) | Aphasic participant 7 | 1 | 79^b^ | 56 | (Semantic judgement>Size judgement) | (fMRI) | (Comprehension) | (High) |
|  |  |  |  |  |  |  |  |  | (Picture naming>Scrambled picture viewing) | (fMRI) | (Production) | (Low) |
| Sebastian 2011 | 10.1080/02687038.2011.557436 | (only one non-aphasic group) | (8) | (unknown) | Aphasic participant 8 | 1 | 60^b^ | 60 | (Semantic judgement>Size judgement) | (fMRI) | (Comprehension) | (High) |
|  |  |  |  |  |  |  |  |  | (Picture naming>Scrambled picture viewing) | (fMRI) | (Production) | (Low) |
| Westmacott 2017 | 10.1017/cjn.2017.44 | (only one non-aphasic group) | 10 | 50.4 | Adult stroke patient 1 | 1 | 45^a^ | 24 | Verb generation>Viewing symbol strings | fMRI | Production | Low |
|  |  |  |  |  |  |  |  |  | Picture-word matching>Symbol matching | fMRI | Comprehension | Low |
| Westmacott 2017 | 10.1017/cjn.2017.44 | (only one non-aphasic group) | (10) | (50.4) | Adult stroke patient 2 | 1 | 49^a^ | 24 | (Verb generation>Viewing symbol strings) | (fMRI) | (Production) | (Low) |
|  |  |  |  |  |  |  |  |  | (Picture-word matching>Symbol matching) | (fMRI) | (Comprehension) | (Low) |
| Westmacott 2017 | 10.1017/cjn.2017.44 | (only one non-aphasic group) | (10) | (50.4) | Adult stroke patient 3 | 1 | 57^a^ | 12 | (Verb generation>Viewing symbol strings) | (fMRI) | (Production) | (Low) |
|  |  |  |  |  |  |  |  |  | (Picture-word matching>Symbol matching) | (fMRI) | (Comprehension) | (Low) |
| Westmacott 2017 | 10.1017/cjn.2017.44 | (only one non-aphasic group) | (10) | (50.4) | Adult stroke patient 4 | 1 | 52^a^ | 24 | (Verb generation>Viewing symbol strings) | (fMRI) | (Production) | (Low) |
|  |  |  |  |  |  |  |  |  | (Picture-word matching>Symbol matching) | (fMRI) | (Comprehension) | (Low) |
| Westmacott 2017 | 10.1017/cjn.2017.44 | (only one non-aphasic group) | (10) | (50.4) | Adult stroke patient 5 | 1 | 43^a^ | 24 | (Verb generation>Viewing symbol strings) | (fMRI) | (Production) | (Low) |
|  |  |  |  |  |  |  |  |  | (Picture-word matching>Symbol matching) | (fMRI) | (Comprehension) | (Low) |
| Abo 2004 | 10.1097/00001756-200408260-00011 | (only one non-aphasic group) | 6 | 21.33 | Aphasic Case 1 | 1 | 56^a^ | 60 | Word repetition>Rest | fMRI | Production | Low |
| Abo 2004 | 10.1097/00001756-200408260-00011 | (only one non-aphasic group) | (6) | (21.33) | Aphasic Case 2 | 1 | 55^a^ | 60 | (Word repetition>Rest) | (fMRI) | (Production) | (Low) |
| Perani 2003 | 10.1016/S0093-934X(02)00561-8 | (only one non-aphasic group) | 10 | (unknown) | Aphasic case 1 | 1 | 46^a^ | 12 | Phonemic fluency>Rest | fMRI | Production | High |
|  |  |  |  |  |  |  |  |  | Semantic fluency>Rest | fMRI | Production | High |
| Perani 2003 | 10.1016/S0093-934X(02)00561-8 | (only one non-aphasic group) | (10) | (unknown) | Aphasic case 2 | 1 | 69^b^ | 12 | (Phonemic fluency>Rest) | (fMRI) | (Production) | (High) |
|  |  |  |  |  |  |  |  |  | (Semantic fluency>Rest) | (fMRI) | (Production) | (High) |
| Perani 2003 | 10.1016/S0093-934X(02)00561-8 | (only one non-aphasic group) | (10) | (unknown) | Aphasic case 3 | 1 | 65^b^ | 10 | (Phonemic fluency>Rest) | (fMRI) | (Production) | (High) |
|  |  |  |  |  |  |  |  |  | (Semantic fluency>Rest) | (fMRI) | (Production) | (High) |
| Perani 2003 | 10.1016/S0093-934X(02)00561-8 | (only one non-aphasic group) | (10) | (unknown) | Aphasic case 4 | 1 | 56^a^ | 24 | (Phonemic fluency>Rest) | (fMRI) | (Production) | (High) |
|  |  |  |  |  |  |  |  |  | (Semantic fluency>Rest) | (fMRI) | (Production) | (High) |
| Rochon 2010 | 10.1016/j.bandl.2010.05.005 | (only one non-aphasic group) | 10 | 61 | Untreated aphasic participant 1 | 1 | 83^b^ | 30 | Semantic judgement>Size judgement | fMRI | Comprehension | High |
|  |  |  |  |  |  |  |  |  | Phonological judgement>Size judgement | fMRI | Production | High |
| Rochon 2010 | 10.1016/j.bandl.2010.05.005 | (only one non-aphasic group) | (10) | (61) | Untreated aphasic participant 2 | 1 | 63^b^ | 48 | (Semantic judgement>Size judgement) | (fMRI) | (Comprehension) | (High) |
|  |  |  |  |  |  |  |  |  | (Phonological judgement>Size judgement) | (fMRI) | (Production) | (High) |
| Sandberg 2014 | 10.1080/13554794.2013.770881 | Normal healthy older adult 1 | 1 | 59.7 | Aphasic participant 1 | 1 | 56^a^ | 38 | Word Judgement abstract words>Control items | fMRI | Comprehension | High |
|  |  |  |  |  |  |  |  |  | Word Judgement concrete words>Control items | fMRI | Comprehension | High |
|  |  |  |  |  |  |  |  |  | Word Judgement abstract words>Concrete words | fMRI | Comprehension | High |
|  |  |  |  |  |  |  |  |  | Word Judgement concrete words>Abstract words | fMRI | Comprehension | High |
|  |  |  |  |  |  |  |  |  | Synonym Judgement abstract words>Control items | fMRI | Comprehension | High |
|  |  |  |  |  |  |  |  |  | Synonym Judgement concrete words>Control items | fMRI | Comprehension | High |
|  |  |  |  |  |  |  |  |  | Synonym Judgement abstract words>Concrete words | fMRI | Comprehension | High |
|  |  |  |  |  |  |  |  |  | Synonym Judgement concrete words>Abstract words | fMRI | Comprehension | High |
| Sandberg 2014 | 10.1080/13554794.2013.770881 | Normal healthy older adult 2 | 1 | (59.7) | Aphasic participant 2 | 1 | 55^a^ | 76 | (Word Judgement abstract words>Control items) | (fMRI) | (Comprehension) | (High) |
|  |  |  |  |  |  |  |  |  | (Word Judgement concrete words>Control items) | (fMRI) | (Comprehension) | (High) |
|  |  |  |  |  |  |  |  |  | (Word Judgement abstract words>Concrete words) | (fMRI) | (Comprehension) | (High) |
|  |  |  |  |  |  |  |  |  | (Word Judgement concrete words>Abstract words) | (fMRI) | (Comprehension) | (High) |
|  |  |  |  |  |  |  |  |  | (Synonym Judgement abstract words>Control items) | (fMRI) | (Comprehension) | (High) |
|  |  |  |  |  |  |  |  |  | (Synonym Judgement concrete words>Control items) | (fMRI) | (Comprehension) | (High) |
|  |  |  |  |  |  |  |  |  | (Synonym Judgement abstract words>Concrete words) | (fMRI) | (Comprehension) | (High) |
|  |  |  |  |  |  |  |  |  | (Synonym Judgement concrete words>Abstract words) | (fMRI) | (Comprehension) | (High) |
| Sandberg 2014 | 10.1080/13554794.2013.770881 | Normal healthy older adult 3 | 1 | (59.7) | Aphasic participant 3 | 1 | 59^b^ | 23 | (Word Judgement abstract words>Control items) | (fMRI) | (Comprehension) | (High) |
|  |  |  |  |  |  |  |  |  | (Word Judgement concrete words>Control items) | (fMRI) | (Comprehension) | (High) |
|  |  |  |  |  |  |  |  |  | (Word Judgement abstract words>Concrete words) | (fMRI) | (Comprehension) | (High) |
|  |  |  |  |  |  |  |  |  | (Word Judgement concrete words>Abstract words) | (fMRI) | (Comprehension) | (High) |
|  |  |  |  |  |  |  |  |  | (Synonym Judgement abstract words>Control items) | (fMRI) | (Comprehension) | (High) |
|  |  |  |  |  |  |  |  |  | (Synonym Judgement concrete words>Control items) | (fMRI) | (Comprehension) | (High) |
|  |  |  |  |  |  |  |  |  | (Synonym Judgement abstract words>Concrete words) | (fMRI) | (Comprehension) | (High) |
|  |  |  |  |  |  |  |  |  | (Synonym Judgement concrete words>Abstract words) | (fMRI) | (Comprehension) | (High) |
| Griffis 2017 | 10.1002/hbm.23476 | (only one non-aphasic group) | 43 | 54 | (only one PSA group) | 43 | 53^a^ | 40.8 | Semantic decision>Tone decision | fMRI | Comprehension | High |
| van Hees 2014 | 10.1016/j.bandl.2013.12.004 | (only one non-aphasic group) | 14 | 61.71 | Aphasic participant 1 | 1 | 60^b^ | 52.3 | Picture naming>Passive viewing of scrambled line drawing | fMRI | Production | Low |
| van Hees 2014 | 10.1016/j.bandl.2013.12.004 | (only one non-aphasic group) | (14) | (61.71) | Aphasic participant 2 | 1 | 60^b^ | (52.3) | (Picture naming>Passive viewing of scrambled line drawing) | (fMRI) | (Production) | (Low) |
| van Hees 2014 | 10.1016/j.bandl.2013.12.004 | (only one non-aphasic group) | (14) | (61.71) | Aphasic participant 3 | 1 | 41^a^ | (52.3) | (Picture naming>Passive viewing of scrambled line drawing) | (fMRI) | (Production) | (Low) |
| van Hees 2014 | 10.1016/j.bandl.2013.12.004 | (only one non-aphasic group) | (14) | (61.71) | Aphasic participant 4 | 1 | 52^a^ | (52.3) | (Picture naming>Passive viewing of scrambled line drawing) | (fMRI) | (Production) | (Low) |
| van Hees 2014 | 10.1016/j.bandl.2013.12.004 | (only one non-aphasic group) | (14) | (61.71) | Aphasic participant 5 | 1 | 56^a^ | (52.3) | (Picture naming>Passive viewing of scrambled line drawing) | (fMRI) | (Production) | (Low) |
| van Hees 2014 | 10.1016/j.bandl.2013.12.004 | (only one non-aphasic group) | (14) | (61.71) | Aphasic participant 6 | 1 | 48^a^ | (52.3) | (Picture naming>Passive viewing of scrambled line drawing) | (fMRI) | (Production) | (Low) |
| van Hees 2014 | 10.1016/j.bandl.2013.12.004 | (only one non-aphasic group) | (14) | (61.71) | Aphasic participant 7 | 1 | 69^b^ | (52.3) | (Picture naming>Passive viewing of scrambled line drawing) | (fMRI) | (Production) | (Low) |
| van Hees 2014 | 10.1016/j.bandl.2013.12.004 | (only one non-aphasic group) | (14) | (61.71) | Aphasic participant 8 | 1 | 65^b^ | (52.3) | (Picture naming>Passive viewing of scrambled line drawing) | (fMRI) | (Production) | (Low) |
| Schofield 2012 | 10.1523/JNEUROSCI.4670-11.2012 | (only one non-aphasic group) | 26 | 54.1 | Moderate comprehension impairment | 12 | 64.8^b^ | 41.2 | Listening to forward or reversed speech>Rest | fMRI | Comprehension | High |
| Schofield 2012 | 10.1523/JNEUROSCI.4670-11.2012 | (only one non-aphasic group) | (26) | (54.1) | Severe comprehension impairment | 9 | 57.0^a^ | 37.4 | (Listening to forward or reversed speech>Rest) | (fMRI) | (Comprehension) | (High) |
| Geranmayeh 2016 | [10.1212/WNL.0000000000002537](https://doi.org/10.1212/WNL.0000000000002537) | (only one non-aphasic group) | 24 | 57 | (only one PSA group) | 53 | 62^b^ | 3.7 | Propositional speech production>Fixation | fMRI | Production | Low |
|  |  |  |  |  |  |  |  |  | Propositional speech production>Counting | fMRI | Production | Low |
|  |  |  |  |  |  |  |  |  | Propositional speech production>Go/no go button press | fMRI | Production | Low |
| Radman 2016 | [10.1155/2016/8797086](https://doi.org/10.1155/2016/8797086) | (only one non-aphasic group) | 5 | 65.6 | (only one PSA group) | 4 | 63.4^b^ | 1.0 (first timepoint) | Picture naming>Fixation | fMRI | Production | Low |
|  |  |  |  |  |  |  |  | 4.3 (last timepoint) |  |  |  |  |
| Long 2017 | 10.1080/02687038.2017.1417538 | (only one non-aphasic group) | 5 | 56.6 | (only one PSA group) | 5 | 55.6^a^ | 0.5-1.2 (first timepoint) | Passive reading>Passive checkerboard viewing | fMRI | Comprehension | Low |
|  |  |  |  |  |  |  |  |  | Picture naming>Scrambled picture viewing | fMRI | Production | Low |
| Long 2017 | 10.1080/02687038.2017.1417538 | (only one non-aphasic group) | (5) | (56.6) | (only one PSA group) | (5) | (55.6^a^) | 11-13 (last timepoint) | (Passive reading>Passive checkerboard viewing) | (fMRI) | (Comprehension) | (Low) |
|  |  |  |  |  |  |  |  |  | (Picture naming>Scrambled picture viewing) | (fMRI) | (Production) | (Low) |
| Hallam 2018 | 10.1016/j.cortex.2017.10.004 | (only one non-aphasic group) | 16 | 64 | (only one PSA group) | 14 | 61^b^ | 85.6 | Passive listening to normal sentences>Spectrally rotated sentences | fMRI | Comprehension | Low |
| Nenert 2018 | 10.3233/RNN-170767 | (only one non-aphasic group) | 85 | 43 | (only one PSA group) | 14 | 46^a^ | 0.5 (first timepoint) | Semantic decision>Tone decision | fMRI | Comprehension | High |
|  |  |  |  |  |  |  |  |  | Covert verb generation>Finger tapping | fMRI | Production | High |
| Nenert 2018 | 10.3233/RNN-170767 | (only one non-aphasic group) | (85) | (43) | (only one PSA group) | (15) | (46^a^) | 3 (last timepoint) | (Semantic decision>Tone decision) | (fMRI) | (Comprehension) | (High) |
|  |  |  |  |  |  |  |  |  | (Covert verb generation>Finger tapping) | (fMRI) | (Production) | (High) |
| Stockert 2020 | 10.1093/brain/awaa023 | (only one non-aphasic group) | 17 | 51.9 | Frontal lesions | 17 | 52.3^a^ | 0.1 (first timepoint) | Passive listening to normal sentences>Reversed sentences | fMRI | Comprehension | High |
| Stockert 2020 | 10.1093/brain/awaa023 | (only one non-aphasic group) | (17) | (51.9) | (Frontal lesions) | (17) | (52.3^a^) | 9.1 (last timepoint) | (Passive listening to normal sentences>Reversed sentences) | (fMRI) | (Comprehension) | (High) |
| Stockert 2020 | 10.1093/brain/awaa023 | (only one non-aphasic group) | (17) | (51.9) | Temporo-parietal lesions | 17 | 54.4^a^ | 0.1 (first timepoint) | (Passive listening to normal sentences>Reversed sentences) | (fMRI) | (Comprehension) | (High) |
| Stockert 2020 | 10.1093/brain/awaa023 | (only one non-aphasic group) | (17) | (51.9) | (Temporo-parietal lesions) | (17) | (54.4^a^) | 8.8 (last timepoint) | (Passive listening to normal sentences>Reversed sentences) | (fMRI) | (Comprehension) | (High) |
| Meier 2019 | 10.1016/j.nicl.2019.101919 | (only one non-aphasic group) | 18 | 59.6 | (only one PSA group) | 34 | 61.9^b^ | 60.0 | Semantic feature judgement>Scrambled picture judgement | fMRI | Comprehension | High |
| Barbieri 2019 | 10.1016/j.cortex.2019.06.015 | (only one non-aphasic group) | 23 | 37.1 | (only one PSA group) | 16 | 48.1^a^ | 49.1 | Picture verification>Scrambled stimuli | fMRI | Comprehension | High |
| Tao 2019 | 10.1016/j.nicl.2019.101865 | (only one non-aphasic group) | 10 | 60.7 | (only one PSA group) | 15 | 61^b^ | 58 | Spelling task>Case verification task | fMRI | Comprehension | High |
| Wilson 2018 | 10.1002/hbm.24077 | (only one non-aphasic group) | 14 | 53.1 | (only one PSA group) | 15 (1 had bilateral strokes) | 60.4^b^ | 65.2 | Narrative comprehension>Backwards speech | fMRI | Comprehension | Low |
|  |  |  |  |  |  |  |  |  | Picture naming>Scrambled picture viewing | fMRI | Production | Low |

Table of all included aphasic groups and their characteristics. Key: a = aphasic group average age ≤57; b = aphasic group average age >57. Abbreviations: ALE = Activation Likelihood Estimation; fMRI = functional Magnetic Resonance Imaging; PET = Positron Emission Tomography; POp = pars opercularis; PSA = Post-stroke aphasia.

Supplementary Table S4: Table of all coordinates included in the omnibus ALE analysis contrasting PSA versus controls performing all tasks

This table is located in a separate Excel file entitled ‘Supplementary File 2’, available through the figshare repository (doi: 10.6084/m9.figshare.12582935).

Table of all coordinates included in the primary ALE contrast analysis comparing all imaging tasks in all PSA vs. all imaging tasks in all controls. The paper, group, imaging contrast and number of participants are provided for all included coordinates at all timepoints at which functional neuroimaging was performed. Abbreviations: PSA = Post-stroke aphasia.

Supplementary Table S5: Significant clusters from the ALE map of all tasks in PSA

| **Single dataset analysis:** | 1521 foci  64 subject groups  481 subjects | |  |  |  |  |
| --- | --- | --- | --- | --- | --- | --- |
| **Cluster** | **Cluster size (mm^3^)** | **x** | **y** | **z** | **Label** | **BA** |
| 1 | 9224 | -46 | 26 | 14 | L Inferior Frontal Gyrus pars triangularis | 45 |
|  |  | -48 | 14 | 16 | L Inferior Frontal Gyrus pars opercularis | 44 |
|  |  | -42 | 30 | -8 | L Frontal Orbital Cortex | 47 |
|  |  | -42 | 10 | 30 | L Middle Frontal Gyrus | 9 |
| 2 | 7464 | 34 | 22 | 0 | R Insular Cortex | * |
|  |  | 44 | 26 | -8 | R Frontal Orbital Cortex | 47 |
|  |  | 46 | 26 | 6 | R Inferior Frontal Gyrus pars triangularis | 13 |
|  |  | 40 | 20 | 22 | R Inferior Frontal Gyrus pars opercularis | 9 |
|  |  | 48 | 26 | 20 | R Inferior Frontal Gyrus pars triangularis | 46 |
|  |  | 46 | 12 | -4 | R Insular Cortex | 13 |
| 3 | 5568 | -6 | 12 | 60 | Superior Frontal Gyrus | 6 |
|  |  | -4 | 2 | 60 | Supplementary Motor Cortex | 6 |
|  |  | -4 | 16 | 42 | Paracingulate Gyrus | 32 |
|  |  | 10 | 16 | 44 | Paracingulate Gyrus | 32 |
|  |  | 8 | 8 | 50 | Supplementary Motor Cortex | 6 |
| 4 | 4984 | 56 | -30 | 4 | R posterior Superior Temporal Gyrus | 22 |
|  |  | 64 | -22 | 2 | R posterior Superior Temporal Gyrus | 41 |
|  |  | 54 | -18 | 10 | R Heschl’s Gyrus (includes H1 and H2) | 41 |
|  |  | 60 | -8 | 0 | R Planum Temporale | * |
| 5 | 2376 | -50 | -38 | 0 | L posterior Middle Temporal Gyrus | 22 |
| 6 | 1616 | 54 | -2 | 40 | R Precentral Gyrus | 6 |
| 7 | 1592 | -46 | -66 | 26 | L superior Lateral Occipital Cortex | 39 |

Table of clusters produced by the ALE single dataset analysis of all imaging tasks in all PSA across all tasks. For each cluster, we provide peak MNI coordinates, cluster size, anatomical label (defined according to the Harvard-Oxford atlas) and Brodmann Area (determined using the Talairach Daemon atlas). The ALE map was thresholded with a voxel-wise uncorrected p<0.001 cluster-forming threshold and a cluster-wise family-wise error corrected threshold of p<0.05 based on 1000 random permutations. Abbreviations: ALE = Activation Likelihood Estimation; L=Left; MNI = Montreal Neurological Institute; R = Right.

Supplementary Table S6: Significant clusters from the ALE map of all tasks in controls

| **Single dataset analysis** | 809 foci  37 subject groups  530 subjects | |  |  |  |  |
| --- | --- | --- | --- | --- | --- | --- |
| **Cluster** | **Cluster size (mm^3^)** | **x** | **y** | **z** | **Label** | **BA** |
| 1 | 12832 | -48 | 12 | 12 | L Inferior Frontal Gyrus pars opercularis | 13 |
|  |  | -38 | 22 | -2 | L Insular Cortex | 13 |
|  |  | -44 | 14 | 26 | L Inferior Frontal Gyrus pars opercularis | 9 |
|  |  | -46 | 30 | 12 | L Inferior Frontal Gyrus pars triangularis | 46 |
|  |  | -48 | 28 | 24 | L Middle Frontal Gyrus | 46 |
|  |  | -44 | 28 | 18 | L Inferior Frontal Gyrus pars triangularis | 46 |
|  |  | -56 | -4 | 24 | L Precentral Gyrus | 6 |
|  |  | -50 | -10 | 30 | L Precentral Gyrus | 6 |
|  |  | -46 | 30 | -2 | L Inferior Frontal Gyrus pars triangularis | 13 |
| 2 | 5992 | 4 | 28 | 38 | Paracingulate Gyrus | 32 |
|  |  | -2 | 26 | 48 | Superior Frontal Gyrus | 8 |
|  |  | -2 | 28 | 40 | Paracingulate Gyrus | 32 |
|  |  | -2 | 12 | 56 | Superior Frontal Gyrus | 6 |
|  |  | -4 | 4 | 60 | Supplementary Motor Cortex | 6 |
|  |  | -6 | 18 | 48 | Paracingulate Gyrus | 6 |
|  |  | 10 | 16 | 46 | Paracingulate Gyrus | 32 |
| 3 | 4080 | -62 | -42 | 4 | L posterior Middle Temporal Gyrus | 22 |
|  |  | -56 | -42 | 4 | L posterior Middle Temporal Gyrus | 22 |
|  |  | -52 | -34 | 8 | L Planum Temporale | 22 |
|  |  | -62 | -30 | 0 | L posterior Superior Temporal Gyrus | 21 |
| 4 | 2816 | 38 | 24 | -6 | R Frontal Orbital Cortex | 13 |
|  |  | 58 | 12 | -14 | R Temporal Pole | 22 |
|  |  | 36 | 32 | 4 | R Inferior Frontal Gyrus pars triangularis | 45 |
|  |  | 46 | 16 | -4 | R Frontal Operculum Cortex | 13 |
| 5 | 1304 | -46 | 4 | 52 | L Middle Frontal Gyrus | 6 |
| 6 | 1240 | -34 | -50 | 44 | L Superior Parietal Lobule | 40 |
|  |  | -40 | -38 | 44 | L Postcentral Gyrus | 40 |
| 7 | 1168 | 64 | -16 | 2 | R posterior Superior Temporal Gyrus | 22 |
|  |  | 66 | -28 | 4 | R posterior Superior Temporal Gyrus | 22 |
|  |  | 66 | -38 | 6 | R posterior Supramarginal Gyrus | 22 |

Table of clusters produced by the ALE single dataset analysis of all imaging tasks in all controls. For each cluster, we provide peak MNI coordinates, cluster size, anatomical label (defined according to the Harvard-Oxford atlas) and Brodmann Area (determined using the Talairach Daemon atlas). The ALE map was thresholded with a voxel-wise uncorrected p<0.001 cluster-forming threshold and a cluster-wise family-wise error corrected threshold of p<0.05 based on 1000 random permutations. Abbreviations: ALE = Activation Likelihood Estimation; L=Left; MNI = Montreal Neurological Institute; R = Right.

Supplementary Table S7: Group similarities between the ALE maps of PSA and controls performing all tasks

| **Conjunction** |  |  |  |  |  |  |
| --- | --- | --- | --- | --- | --- | --- |
| **Cluster** | **Cluster size (mm^3^)** | **x** | **y** | **z** | **Label** | **BA** |
| 1 | 6088 | -46 | 30 | 12 | L Inferior Frontal Gyrus pars triangularis | 46 |
|  |  | -44 | 28 | 18 | L Inferior Frontal Gyrus pars triangularis | 46 |
|  |  | -48 | 14 | 16 | L Inferior Frontal Gyrus pars opercularis | 44 |
|  |  | -44 | 30 | -4 | L Frontal Orbital Cortex | 47 |
|  |  | -36 | 28 | -8 | L Frontal Orbital Cortex | 47 |
|  |  | -42 | 20 | 2 | L Frontal Operculum Cortex | 13 |
|  |  | -42 | 10 | 30 | L Middle Frontal Gyrus | 9 |
| 2 | 1544 | -54 | -40 | 2 | L posterior Middle Temporal Gyrus | 22 |
| 3 | 1176 | 38 | 24 | -4 | R Frontal Orbital Cortex | 13 |
|  |  | 48 | 18 | -4 | R Frontal Operculum Cortex | 13 |
|  |  | 38 | 26 | 2 | R Frontal Operculum Cortex | 13 |
| 4 | 824 | 64 | -16 | 2 | R posterior Superior Temporal Gyrus | 22 |
|  |  | 66 | -26 | 2 | R posterior Superior Temporal Gyrus | 22 |
| 5 | 584 | -4 | 14 | 56 | Superior Frontal Gyrus | 6 |
|  |  | -6 | 16 | 48 | Paracingulate gyrus | 32 |
| 6 | 464 | -4 | 4 | 60 | Supplementary Motor Cortex | 6 |
| 7 | 304 | -4 | 12 | 56 | Superior Frontal Gyrus | 6 |
| 8 | 200 | 10 | 16 | 46 | Paracingulate gyrus | 32 |
| 9 | 184 | -4 | 8 | 60 | Supplementary Motor Cortex | 6 |
| 10 | 120 | 6 | 22 | 44 | Paracingulate gyrus | 6 |
|  |  | -2 | 22 | 42 | Paracingulate gyrus | 32 |
| 11 | 104 | -6 | 4 | 60 | Supplementary Motor Cortex | 6 |
| 12 | 96 | 8 | 18 | 46 | Paracingulate gyrus | 6 |
| 13 | 88 | -8 | 6 | 62 | Supplementary Motor Cortex | 6 |
| 14 | 72 | -8 | 16 | 50 | Superior Frontal Gyrus | 6 |
| 15 | 64 | -6 | 12 | 54 | Superior Frontal Gyrus | 6 |
| 16 | 64 | -6 | 10 | 60 | Superior Frontal Gyrus | 6 |
| 17 | 48 | -6 | 6 | 60 | Supplementary Motor Cortex | 6 |
| 18 | 40 | -8 | 4 | 62 | Supplementary Motor Cortex | 6 |
| 19 | 32 | 6 | 20 | 44 | Paracingulate Gyrus | 6 |
| 20 | 32 | -8 | 14 | 50 | Paracingulate Gyrus | 6 |
| 21 | 24 | -4 | -2 | 60 | Supplementary Motor Cortex | 6 |
| 22 | 16 | 6 | 22 | 42 | Paracingulate Gyrus | 6 |
| 23 | 16 | 8 | 14 | 44 | Paracingulate Gyrus | 32 |
| 24 | 8 | 62 | -36 | 6 | R posterior Supramarginal Gyrus | 22 |
| 25 | 25 | 8 | 20 | 42 | Paracingulate Gyrus | 32 |
| 26 | 8 | -8 | 12 | 50 | Paracingulate Gyrus | 6 |

Table of clusters produced by the primary ALE contrast analysis showing the conjunction between all imaging tasks in all PSA with all imaging tasks in all controls. For each cluster, we provide peak MNI coordinates, cluster size, anatomical label (defined according to the Harvard-Oxford atlas) and Brodmann Area (determined using the Talairach Daemon atlas). Abbreviations: ALE = Activation Likelihood Estimation.

Supplementary Table S8: Group differences between the ALE maps of PSA versus controls performing all tasks

| **Cluster** | **Cluster size (mm^3^)** | **x** | **y** | **z** | **Z (peak)** | **Label** | **BA** |
| --- | --- | --- | --- | --- | --- | --- | --- |
| **PSA>Controls** |  |  |  |  |  |  |  |
| 1 | 280 | 40 | 20 | 12 | 2.32 | R Inferior Frontal Gyrus pars opercularis | 13 |
|  |  | 40 | 18 | 6 | 2.05 | R Frontal Operculum | 13 |
| **Controls>PSA** |  |  |  |  |  |  |  |
| 1 | 4024 | -50 | 6 | 24 | 3.06 | L Precentral Gyrus | 6 |
|  |  | -52 | 12 | 10 | 2.99 | L Inferior Frontal Gyrus pars opercularis | 44 |
|  |  | -56 | 26 | 12 | 2.24 | L Inferior Frontal Gyrus pars triangularis | 45 |
|  |  | -58 | -8 | 22 | 2.16 | L Postcentral Gyrus | 4 |
|  |  | -62 | -4 | 22 | 2.15 | L Precentral Gyrus | 4 |
|  |  | -52 | -12 | 30 | 1.91 | L Postcentral Gyrus | 6 |
|  |  | -42 | 14 | 32 | 1.70 | L Middle Frontal Gyrus | 9 |
| 2 | 2504 | -2 | 28 | 46 | 2.70 | Superior Frontal Gyrus | 8 |
|  |  | -2 | 24 | 48 | 2.56 | Superior Frontal Gyrus | 8 |
|  |  | -4 | 30 | 34 | 2.37 | Paracingulate Gyrus | 32 |
|  |  | 6 | 28 | 46 | 2.27 | Superior Frontal Gyrus | 8 |
|  |  | 6 | 26 | 40 | 2.21 | Paracingulate Gyrus | 32 |
|  |  | -6 | 2 | 52 | 2.18 | Supplementary Motor Cortex | 6 |
|  |  | 2 | 16 | 54 | 2.09 | Superior Frontal Gyrus | 6 |
|  |  | -2 | 8 | 54 | 2.04 | Supplementary Motor Cortex | 6 |
| 3 | 1968 | -50 | -34 | 12 | 2.82 | L Planum Temporale | 41 |
|  |  | -50 | -30 | 10 | 2.67 | L Planum Temporale | 41 |
|  |  | -60 | -48 | 4 | 2.64 | L temporooccipital Middle Temporal Gyrus | 22 |
| 4 | 1240 | -34 | -44 | 44 | 3.89 | L Superior Parietal Lobule | 40 |
|  |  | -30 | -54 | 46 | 3.43 | L Superior Parietal Lobule | 7 |
|  |  | -40 | -36 | 42 | 2.47 | L Postcentral Gyrus | 40 |
| 5 | 1152 | -36 | 26 | 0 | 2.89 | L Frontal Orbital Cortex | 13 |
| 6 | 920 | -44 | 6 | 56 | 2.97 | L Middle Frontal Gyrus | 6 |
| 7 | 592 | 58 | 14 | -10 | 2.83 | R Temporal Pole | 22 |
| 8 | 528 | -50 | 31 | 28 | 2.51 | L Middle Frontal Gyrus | 9 |
| 9 | 496 | 34 | 34 | 4 | 2.44 | R Inferior Frontal Gyrus pars triangularis | 45 |

Table of clusters produced by the primary ALE contrast analysis showing differences between all imaging tasks in all PSA vs. all imaging tasks in all controls. For each cluster, we provide the peak MNI coordinate, cluster size, anatomical label (defined according to the Harvard-Oxford atlas) and Brodmann Area (determined using the Talairach Daemon atlas). Thresholded ALE maps from the two datasets being compared were subtracted from each other and thresholded at p<0.05 using 10000 P-value permutations with a minimum cluster threshold of 200mm^3^. Abbreviations: ALE = Activation Likelihood Estimation; BA = Brodmann Area; L=Left; MNI = Montreal Neurological Institute; R = Right. PSA = Post-stroke aphasia.

Supplementary Table S9: Significant clusters from the ALE map of PSA performing comprehension tasks

| **Single dataset analysis** | 1018 foci  40 subject groups  306 subjects | |  |  |  |  |
| --- | --- | --- | --- | --- | --- | --- |
| **Cluster** | **Cluster size (mm^3^)** | **x** | **y** | **z** | **Label** | **BA** |
| 1 | 6216 | -46 | 24 | 16 | L Inferior Frontal Gyrus pars triangularis | 46 |
|  |  | -46 | 14 | 16 | L Inferior Frontal Gyrus pars opercularis | 13 |
|  |  | -38 | 30 | -10 | L Frontal Orbital Cortex | 47 |
| 2 | 3640 | 34 | 22 | -2 | R Insular Cortex | * |
|  |  | 44 | 26 | -8 | R Frontal Orbital Cortex | 47 |
| 3 | 1896 | -52 | -38 | 0 | L posterior Middle Temporal Gyrus | 22 |
| 4 | 1664 | -42 | -64 | 26 | L superior Lateral Occipital Cortex | 39 |
| 5 | 1424 | -6 | 16 | 52 | Superior Frontal Gyrus | 6 |
|  |  | -6 | 12 | 60 | Superior Frontal Gyrus | 6 |
|  |  | -2 | 0 | 60 | Supplementary Motor Cortex | 6 |
| 6 | 1160 | 40 | 22 | 22 | R Middle Frontal Gyrus | 9 |
|  |  | 48 | 26 | 20 | R Inferior Frontal Gyrus pars triangularis | 46 |

Table of clusters produced by the ALE single dataset analysis of comprehension tasks in PSA. For each cluster, we provide peak MNI coordinates, cluster size, anatomical label (defined according to the Harvard-Oxford atlas) and Brodmann Area (determined using the Talairach Daemon atlas). The ALE map was thresholded with a voxel-wise uncorrected p<0.001 cluster-forming threshold and a cluster-wise family-wise error corrected threshold of p<0.05 based on 1000 random permutations. Abbreviations: ALE = Activation Likelihood Estimation; BA = Brodmann Area; L=Left; MNI = Montreal Neurological Institute; R = Right. PSA = Post-stroke aphasia.

Supplementary Table S10: Significant clusters from the ALE map of controls performing comprehension tasks

| **Single dataset analysis** | 388 foci  25 subject groups  358 subjects | |  |  |  |  |
| --- | --- | --- | --- | --- | --- | --- |
| **Cluster** | **Cluster size (mm^3^)** | **x** | **y** | **z** | **Label** | **BA** |
| 1 | 3968 | -42 | 12 | 28 | L Inferior Frontal Gyrus pars opercularis | 9 |
|  |  | -48 | 14 | 14 | L Inferior Frontal Gyrus pars opercularis | 44 |
|  |  | -46 | 28 | 26 | L Middle Frontal Gyrus | 9 |
|  |  | -46 | 26 | 16 | L Inferior Frontal Gyrus pars triangularis | 46 |
|  |  | -44 | 24 | 30 | L Middle Frontal Gyrus | 9 |
| 2 | 1832 | -46 | 32 | -4 | L Inferior Frontal Gyrus pars triangularis | 47 |
|  |  | -38 | 30 | -8 | L Frontal Orbital Cortex | 47 |
|  |  | -44 | 28 | -20 | L Frontal Orbital Cortex | 47 |
| 3 | 1152 | -4 | 48 | 34 | Superior Frontal Gyrus | 6 |
|  |  | -6 | 54 | 24 | Superior Frontal Gyrus | 9 |
|  |  | 0 | 44 | 24 | Paracingulate Gyrus | 9 |
| 4 | 1104 | -48 | -52 | -10 | L temporooccipital Inferior Temporal Gyrus | 37 |
|  |  | -46 | -50 | -16 | L temporooccipital Inferior Temporal Gyrus | 37 |
| 5 | 1088 | -56 | -40 | 2 | L posterior Middle Temporal Gyrus | 22 |
| 6 | 1024 | -46 | 4 | 52 | L Middle Frontal Gyrus | 6 |

Table of clusters produced by the ALE single dataset analysis of comprehension tasks in controls. For each cluster, we provide peak MNI coordinates, cluster size, anatomical label (defined according to the Harvard-Oxford atlas) and Brodmann Area (determined using the Talairach Daemon atlas). The ALE map was thresholded with a voxel-wise uncorrected p<0.001 cluster-forming threshold and a cluster-wise family-wise error corrected threshold of p<0.05 based on 1000 random permutations. Abbreviations: ALE = Activation Likelihood Estimation; BA = Brodmann Area; L=Left; MNI = Montreal Neurological Institute; R = Right.

Supplementary Table S11: Group similarities between the ALE maps of PSA and controls performing comprehension tasks

| **Conjunction** |  |  |  |  |  |  |
| --- | --- | --- | --- | --- | --- | --- |
| **Cluster** | **Cluster size (mm^3^)** | **x** | **y** | **z** | **Label** | **BA** |
| 1 | 2064 | -48 | 14 | 16 | L Inferior Frontal Gyrus pars opercularis | 44 |
|  |  | -46 | 26 | 16 | L Inferior Frontal Gyrus pars triangularis | 46 |
|  |  | -46 | 28 | 22 | L Inferior Frontal Gyrus pars triangularis | 46 |
|  |  | -44 | 14 | 22 | L Inferior Frontal Gyrus pars opercularis | 9 |
| 2 | 720 | -56 | -42 | 2 | L posterior Middle Temporal Gyrus | 22 |
| 3 | 560 | -36 | 28 | -8 | L Frontal Orbital Cortex | 47 |
|  |  | -40 | 30 | -6 | L Frontal Orbital Cortex | 47 |

Table of clusters produced by the ALE contrast analysis showing the conjunction between comprehension tasks in PSA with comprehension tasks in controls. For each cluster, we provide peak MNI coordinates, cluster size, anatomical label (defined according to the Harvard-Oxford atlas) and Brodmann Area (determined using the Talairach Daemon atlas). Abbreviations: ALE = Activation Likelihood Estimation; L=Left; MNI = Montreal Neurological Institute; R = Right.

Supplementary Table S12: Group differences between the ALE maps of PSA versus controls performing comprehension tasks

| **Cluster** | **Cluster size (mm^3^)** | **x** | **y** | **z** | **Z (peak)** | **Label** | **BA** |
| --- | --- | --- | --- | --- | --- | --- | --- |
| **PSA>Controls (comprehension)** |  |  |  |  |  |  |  |
| 1 | 424 | 34 | 18 | 4 | 2.20 | R insular cortex | * |
| **Controls>PSA (comprehension)** |  |  |  |  |  |  |  |
| 1 | 712 | -4 | 48 | 26 | 2.31 | Paracingulate Gyrus | 9 |
|  |  | -6 | 54 | 28 | 2.21 | Superior Frontal Gyrus | 9 |
|  |  | -4 | 54 | 20 | 2.06 | Superior Frontal Gyrus | 9 |
| 2 | 576 | -44 | 4 | 52 | 2.25 | L Middle Frontal Gyrus | 6 |
| 3 | 352 | -46 | 16 | 30 | 2.42 | L Middle Frontal Gyrus | 9 |
| 4 | 232 | -52 | 32 | -4 | 2.24 | L Inferior Frontal Gyrus pars triangularis | 47 |

Table of clusters produced by the ALE contrast analysis showing differences between comprehension tasks in PSA vs. comprehension tasks in controls. For each cluster, we provide the peak MNI coordinate, cluster size, anatomical label (defined according to the Harvard-Oxford atlas) and Brodmann Area (determined using the Talairach Daemon atlas). Thresholded ALE maps from the two datasets being compared were subtracted from each other and thresholded at p<0.05 using 10000 P-value permutations with a minimum cluster threshold of 200mm^3^. Abbreviations: ALE = Activation Likelihood Estimation; L=Left; MNI = Montreal Neurological Institute; R = Right. PSA = Post-stroke aphasia.

Supplementary Table S13: Significant clusters from the ALE map of PSA performing production tasks

| **Single dataset analysis** | 503 foci  42 subject groups  225 subjects | |  |  |  |  |
| --- | --- | --- | --- | --- | --- | --- |
| **Cluster** | **Cluster size (mm^3^)** | **x** | **y** | **z** | **Label** | **BA** |
| 1 | 5176 | 56 | -30 | 4 | R posterior Superior Temporal Gyrus | 22 |
|  |  | 64 | -20 | 0 | R posterior Superior Temporal Gyrus | * |
|  |  | 54 | -18 | 10 | R Heschl’s Gyrus (includes H1 and H2) | 41 |
|  |  | 44 | -22 | 12 | R Heschl’s Gyrus (includes H1 and H2) | 41 |
| 2 | 2464 | -46 | 32 | 14 | L Inferior Frontal Gyrus pars triangularis | 46 |
|  |  | -54 | 24 | -2 | L Inferior Frontal Gyrus pars triangularis | 47 |
| 3 | 1920 | 50 | -4 | 38 | R Precentral Gyrus | 6 |
|  |  | 54 | 4 | 36 | R Precentral Gyrus | 6 |
| 4 | 1232 | 50 | 20 | -4 | R Inferior Frontal Gyrus pars triangularis | 47 |
|  |  | 46 | 12 | -4 | R Insular Cortex | 13 |
| 5 | 1072 | 8 | 8 | 50 | Supplementary Motor Cortex | 6 |
|  |  | 12 | 18 | 42 | R Paracingulate Gyrus | 32 |
| 6 | 808 | -6 | 12 | 60 | Superior Frontal Gyrus | 6 |
|  |  | -4 | -2 | 66 | Supplementary Motor Cortex | 6 |

Table of clusters produced by the ALE single dataset analysis of production tasks in PSA. For each cluster, we provide peak MNI coordinates, cluster size, anatomical label (defined according to the Harvard-Oxford atlas) and Brodmann Area (determined using the Talairach Daemon atlas). The ALE map was thresholded with a voxel-wise uncorrected p<0.001 cluster-forming threshold and a cluster-wise family-wise error corrected threshold of p<0.05 based on 1000 random permutations. Abbreviations: ALE = Activation Likelihood Estimation; L=Left; MNI = Montreal Neurological Institute; R = Right.

Supplementary Table S14: Significant clusters from the ALE map of controls performing production tasks

| **Single dataset analysis** | 421 foci  18 subject groups  304 subjects | |  |  |  |  |
| --- | --- | --- | --- | --- | --- | --- |
| **Cluster** | **Cluster size (mm^3^)** | **x** | **y** | **z** | **Label** | **BA** |
| 1 | 4952 | -2 | 28 | 38 | Paracingulate Gyrus | 32 |
|  |  | -4 | 2 | 60 | Supplementary Motor Cortex | 6 |
|  |  | 4 | 28 | 38 | Paracingulate Gyrus | 32 |
|  |  | 10 | 16 | 46 | Paracingulate Gyrus | 32 |
|  |  | 0 | 26 | 48 | Superior Frontal Gyrus | 8 |
|  |  | -4 | 12 | 52 | Paracingulate Gyrus | 6 |
|  |  | -4 | 2 | 70 | Supplementary Motor Cortex | 6 |
| 2 | 4680 | -56 | -4 | 24 | L Precentral Gyrus | 6 |
|  |  | -50 | -10 | 30 | L Precentral Gyrus | 6 |
|  |  | -50 | 30 | 10 | L Inferior Frontal Gyrus pars triangularis | 46 |
|  |  | -50 | 12 | 10 | L Inferior Frontal Gyrus pars opercularis | 44 |
|  |  | -54 | 16 | 6 | L Inferior Frontal Gyrus pars opercularis | 44 |
|  |  | -52 | 20 | 8 | L Inferior Frontal Gyrus pars opercularis | 44 |
|  |  | -50 | 28 | 24 | L Middle Frontal Gyrus | 46 |
|  |  | -48 | -12 | 40 | L Precentral Gyrus | 4 |
|  |  | -42 | 28 | 18 | L Inferior Frontal Gyrus pars triangularis | 46 |
|  |  | -48 | 8 | 20 | L Inferior Frontal Gyrus pars opercularis | 9 |
| 3 | 2560 | -52 | -34 | 8 | L Planum Temporale | 22 |
|  |  | -62 | -42 | 6 | L posterior Supramarginal Gyrus | 22 |
|  |  | -56 | -46 | 6 | L temporooccipital Middle Temporal Gyrus | 22 |
|  |  | -50 | -40 | -2 | L posterior Middle Temporal Gyrus | 22 |
| 4 | 1264 | 68 | -28 | 2 | R posterior Superior Temporal Gyrus | 22 |
| 5 | 1080 | -38 | 22 | -2 | L Insular Cortex | 13 |

Table of clusters produced by the ALE single dataset analysis of production tasks in controls. For each cluster, we provide peak MNI coordinates, cluster size, anatomical label (defined according to the Harvard-Oxford atlas) and Brodmann Area (determined using the Talairach Daemon atlas). The ALE map was thresholded with a voxel-wise uncorrected p<0.001 cluster-forming threshold and a cluster-wise family-wise error corrected threshold of p<0.05 based on 1000 random permutations. Abbreviations: ALE = Activation Likelihood Estimation; L=Left; MNI = Montreal Neurological Institute; R = Right.

Supplementary Table S15: Group similarities between the ALE maps of PSA and controls performing production tasks

| **Conjunction** |  |  |  |  |  |  |
| --- | --- | --- | --- | --- | --- | --- |
| **Cluster** | **Cluster size (mm^3^)** | **x** | **y** | **z** | **Label** | **BA** |
| 1 | 680 | -48 | 30 | 12 | L Inferior Frontal Gyrus pars triangularis | 46 |
| 2 | 656 | 66 | -22 | 2 | R posterior Superior Temporal Gyrus | * |
|  |  | 62 | -30 | 4 | R posterior Superior Temporal Gyrus | 22 |
| 3 | 376 | 10 | 16 | 44 | Paracingulate Gyrus | 32 |
| 4 | 312 | -4 | 12 | 58 | Superior Frontal Gyrus | 6 |
|  |  | -4 | 8 | 60 | Supplementary Motor Cortex | 6 |
|  |  | -4 | 0 | 64 | Supplementary Motor Cortex | 6 |
| 5 | 8 | 66 | -32 | 2 | R posterior Superior Temporal Gyrus | 22 |

Table of clusters produced by the ALE contrast analysis showing the conjunction between production tasks in PSA with production tasks in controls. For each cluster, we provide peak MNI coordinates, cluster size, anatomical label (defined according to the Harvard-Oxford atlas) and Brodmann Area (determined using the Talairach Daemon atlas). Abbreviations: ALE = Activation Likelihood Estimation; BA = Brodmann Area; L=Left; MNI = Montreal Neurological Institute; R = Right. PSA = Post-stroke aphasia.

Supplementary Table S16: Group differences between the ALE maps of PSA versus controls performing production tasks

| **Cluster** | **Cluster size (mm^3^)** | **x** | **y** | **z** | **Z (peak)** | **Label** | **BA** |
| --- | --- | --- | --- | --- | --- | --- | --- |
| **PSA>Controls (production)** |  |  |  |  |  |  |  |
| No clusters found |  |  |  |  |  |  |  |
| **Controls>PSA (production)** |  |  |  |  |  |  |  |
| 1 | 6040 | -54 | 0 | 22 | 3.16 | L Precentral Gyrus | 6 |
|  |  | -36 | 26 | 2 | 2.97 | L Frontal Operculum Cortex | 13 |
|  |  | -42 | 22 | -4 | 2.75 | L Frontal Orbital Cortex | 13 |
|  |  | -52 | 10 | 12 | 2.73 | L Inferior Frontal Gyrus pars opercularis | 44 |
|  |  | -62 | -4 | 22 | 2.64 | L Precentral Gyrus | 4 |
|  |  | -56 | 18 | 10 | 2.62 | L Inferior Frontal Gyrus pars opercularis | 44 |
|  |  | -54 | 26 | 12 | 2.54 | L Inferior Frontal Gyrus pars triangularis | 45 |
|  |  | -54 | -12 | 30 | 2.44 | L Postcentral Gyrus | 4 |
|  |  | -50 | 8 | 22 | 2.37 | L Precentral Gyrus | 9 |
|  |  | -46 | 28 | 22 | 2.25 | L Inferior Frontal Gyrus pars triangularis | 46 |
| 2 | 4496 | 4 | 26 | 40 | 3.29 | Paracingulate Gyrus | 32 |
|  |  | 6 | 18 | 54 | 2.89 | Superior Frontal Gyrus | 6 |
|  |  | 2 | 22 | 50 | 2.70 | Superior Frontal Gyrus | 6 |
|  |  | -2 | 16 | 52 | 2.53 | Superior Frontal Gyrus | 6 |
|  |  | 0 | 8 | 56 | 2.35 | Supplementary Motor Cortex | 6 |
|  |  | -2 | 4 | 62 | 2.29 | Supplementary Motor Cortex | 6 |
|  |  | -6 | 2 | 74 | 2.12 | Superior Frontal Gyrus | 6 |
|  |  | -6 | 2 | 54 | 2.06 | Supplementary Motor Cortex | 6 |
|  |  | 14 | 16 | 50 | 1.81 | R Superior Frontal Gyrus | 6 |
| 3 | 2888 | -48 | -30 | 10 | 3.72 | L Planum Temporale | 41 |
|  |  | -56 | -40 | 4 | 3.54 | L posterior Superior Temporal Gyrus | 22 |
| 4 | 952 | 68 | -30 | 8 | 2.81 | R posterior Superior Temporal Gyrus | 42 |
| 5 | 560 | 36 | 26 | -2 | 2.40 | R Frontal Orbital Cortex | 13 |
|  |  | 36 | 28 | -6 | 2.38 | R Frontal Orbital Cortex | 47 |
|  |  | 38 | 20 | -6 | 2.13 | R Insular Cortex | 47 |
| 6 | 488 | 48 | -2 | 28 | 2.54 | R Precentral Gyrus | 6 |
|  |  | 54 | -4 | 26 | 2.37 | R Precentral Gyrus | 6 |
| 7 | 392 | 56 | 16 | -10 | 2.64 | R Temporal Pole | 22 |
| 8 | 264 | -68 | -22 | 2 | 2.27 | L posterior Superior Temporal Gyrus | 22 |
| 9 | 200 | 52 | -12 | 2 | 1.95 | R Heschl’s Gyrus (includes H1 and H2) | 22 |

Table of clusters produced by the ALE contrast analysis showing differences between production tasks in PSA vs. production tasks in controls. For each cluster, we provide the peak MNI coordinate, cluster size, anatomical label (defined according to the Harvard-Oxford atlas) and Brodmann Area (determined using the Talairach Daemon atlas). Thresholded ALE maps from the two datasets being compared were subtracted from each other and thresholded at p<0.05 using 10000 P-value permutations with a minimum cluster threshold of 200mm^3^. Abbreviations: ALE = Activation Likelihood Estimation; BA = Brodmann Area; L=Left; MNI = Montreal Neurological Institute; R = Right. PSA = Person(s) with Post-stroke aphasia. * = No Brodmann Area according to Talairach Daemon.

Supplementary Table S17: Group differences between the ALE maps of PSA performing comprehension tasks versus production tasks

| **Cluster** | **Cluster size (mm^3^)** | **x** | **y** | **z** | **Z (peak)** | **Label** | **BA** |
| --- | --- | --- | --- | --- | --- | --- | --- |
| **Comprehension>Production (PSA)** |  |  |  |  |  |  |  |
| 1 | 2128 | -44 | 18 | 16 | 3.29 | L Inferior Frontal Gyrus pars opercularis | 46 |
| 2 | 1976 | 28 | 22 | 2 | 2.75 | R Insular Cortex | * |
|  |  | 36 | 20 | -8 | 2.73 | R Insular Cortex | 47 |
| 3 | 1080 | -40 | -68 | 20 | 3.09 | L superior Lateral Occipital Cortex | 39 |
|  |  | -36 | -64 | 26 | 2.93 | L Angular Gyrus | 19 |
| 4 | 1040 | -56 | -39 | -6 | 2.73 | L posterior Middle Temporal Gyrus | * |
| 5 | 1008 | -32 | 28 | -12 | 2.69 | L Frontal Orbital Cortex | 47 |
|  |  | -36 | 28 | -14 | 2.66 | L Frontal Orbital Cortex | 47 |
| 6 | 384 | 44 | 28 | 24 | 2.13 | R Middle Frontal Gyrus | 9 |
| **Production> Comprehension (PSA)** |  |  |  |  |  |  |  |
| 1 | 2216 | 60 | -32 | 8 | 2.81 | R posterior Superior Temporal Gyrus | 41 |
|  |  | 54 | -26 | 8 | 2.49 | R Planum Temporale | 41 |
| 2 | 496 | 48 | -10 | 36 | 2.43 | R Precentral Gyrus | 6 |
|  |  | 52 | -6 | 32 | 2.14 | R Precentral Gyrus | 6 |
| 3 | 224 | 51 | 16 | 0 | 2.05 | R Inferior Frontal Gyrus pars opercularis | 13 |

Table of clusters produced by the ALE contrast analysis showing differences between PSA performing comprehension tasks versus PSA performing production tasks. For each cluster, we provide the peak MNI coordinate, cluster size, anatomical label (defined according to the Harvard-Oxford atlas) and Brodmann Area (determined using the Talairach Daemon atlas). Thresholded ALE maps from the two datasets being compared were subtracted from each other and thresholded at p<0.05 using 10000 P-value permutations with a minimum cluster threshold of 200mm^3^. Abbreviations: ALE = Activation Likelihood Estimation; BA = Brodmann Area; L=Left; MNI = Montreal Neurological Institute; R = Right. PSA = Person(s) with Post-stroke aphasia. * = No Brodmann Area according to Talairach Daemon.

Supplementary Table S18: Group differences between the ALE maps of controls performing comprehension tasks versus production tasks

| **Cluster** | **Cluster size (mm^3^)** | **x** | **y** | **z** | **Z (peak)** | **Label** | **BA** |
| --- | --- | --- | --- | --- | --- | --- | --- |
| **Comprehension>Production (controls)** |  |  |  |  |  |  |  |
| 1 | 1096 | -44 | 32 | -22 | 3.54 | L Frontal Pole | 47 |
|  |  | -44 | 32 | -16 | 3.43 | L Frontal Orbital Cortex | 47 |
|  |  | -43 | 27 | -22 | 3.35 | L Temporal Pole | 47 |
|  |  | -40 | 32 | -18 | 3.12 | L Frontal Orbital Cortex | 47 |
|  |  | -46 | 34 | -8 | 3.01 | L Frontal Orbital Cortex | 47 |
| 2 | 896 | 0 | 43 | 23 | 3.35 | Paracingulate Gyrus | 9 |
|  |  | 0 | 44 | 28 | 3.29 | Paracingulate Gyrus | 9 |
|  |  | -2 | 44 | 36 | 2.65 | Superior Frontal Gyrus | 8 |
| 3 | 568 | -48 | -46 | -18 | 2.40 | L temporooccipital Inferior Temporal Gyrus | 37 |
| **Production> Comprehension (controls)** |  |  |  |  |  |  |  |
| 1 | 2096 | -56 | -6 | 26 | 3.54 | L Precentral Gyrus | 4 |
|  |  | -51 | -6 | 29 | 3.16 | L Precentral Gyrus | 6 |
|  |  | -52 | 2 | 24 | 2.97 | L Precentral Gyrus | 9 |
|  |  | -48 | -10 | 40 | 2.35 | L Precentral Gyrus | 4 |
| 2 | 1792 | 8 | 20 | 48 | 2.75 | Paracingulate Gyrus | 6 |
|  |  | 6 | 18 | 52 | 2.67 | Superior Frontal Gyrus | 6 |
|  |  | 0 | 24 | 40 | 2.21 | Paracingulate Gyrus | 32 |
| 3 | 1264 | -48 | -30 | 10 | 3.12 | L Planum Temporale | 41 |
|  |  | -52 | -42 | 12 | 2.52 | L posterior Supramarginal Gyrus | 22 |
|  |  | -52 | -48 | 6 | 1.89 | L temporooccipital Middle Temporal Gyrus | 21 |
| 4 | 1120 | 66 | -27 | 0 | 3.89 | R posterior Superior Temporal Gyrus | * |
|  |  | 68 | -26 | 6 | 3.72 | R posterior Superior Temporal Gyrus | 22 |
| 5 | 1096 | -2 | 1 | 67 | 3.09 | Supplementary Motor Cortex | 6 |
|  |  | -4 | 4 | 69 | 2.70 | Supplementary Motor Cortex | 6 |
| 6 | 440 | -50 | 20 | 2 | 2.09 | L Inferior Frontal Gyrus pars triangularis | 44 |
|  |  | -56 | 18 | 10 | 1.88 | L Inferior Frontal Gyrus pars opercularis | 44 |
|  |  | -52 | 26 | 10 | 1.83 | L Inferior Frontal Gyrus pars triangularis | 45 |
| 7 | 336 | -36 | 28 | 2 | 2.18 | L Frontal Orbital Cortex | 45 |
|  |  | -40 | 18 | -4 | 1.83 | L Insular Cortex | * |
| 8 | 232 | 52 | 18 | -8 | 2.51 | R Temporal Pole | 47 |

Table of clusters produced by the ALE contrast analysis showing differences between controls performing comprehension tasks versus controls performing production tasks. For each cluster, we provide the peak MNI coordinate, cluster size, anatomical label (defined according to the Harvard-Oxford atlas) and Brodmann Area (determined using the Talairach Daemon atlas). Thresholded ALE maps from the two datasets being compared were subtracted from each other and thresholded at p<0.05 using 10000 P-value permutations with a minimum cluster threshold of 200mm^3^. Abbreviations: ALE = Activation Likelihood Estimation; BA = Brodmann Area; L=Left; MNI = Montreal Neurological Institute; R = Right. PSA = Person(s) with Post-stroke aphasia. * = No Brodmann Area according to Talairach Daemon.

Supplementary Table S19: Significant clusters from the ALE map of PSA performing higher demand comprehension tasks

| **Single dataset analysis** | 840 foci  28 subject groups  230 subjects | |  |  |  |  |
| --- | --- | --- | --- | --- | --- | --- |
| **Cluster** | **Cluster size (mm^3^)** | **x** | **y** | **z** | **Label** | **BA** |
| 1 | 4904 | 34 | 22 | -2 | R Insular Cortex | * |
|  |  | 44 | 26 | -8 | R Frontal Orbital Cortex | 47 |
| 2 | 4880 | -46 | 24 | 16 | L Inferior Frontal Gyrus pars triangularis | 46 |
| 3 | 2056 | -42 | -64 | 26 | L superior Lateral Occipital Cortex | 39 |
| 4 | 1696 | -6 | 16 | 52 | Superior Frontal Gyrus | 6 |
|  |  | -6 | 12 | 60 | Superior Frontal Gyrus | 6 |
|  |  | -2 | 0 | 60 | Supplementary Motor Cortex | 6 |
| 5 | 1576 | 40 | 22 | 22 | R Middle Frontal Gyrus | 9 |
|  |  | 48 | 28 | 20 | R Inferior Frontal Gyrus pars triangularis | 46 |
| 6 | 904 | -34 | 34 | -14 | L Frontal Orbital Cortex | 47 |
|  |  | -42 | 30 | -4 | L Frontal Orbital Cortex | 47 |
|  |  | -36 | 30 | -10 | L Frontal Orbital Cortex | 47 |
| 7 | 816 | -54 | -10 | -12 | L anterior Superior Temporal Gyrus | 22 |

Table of clusters produced by the ALE single dataset analysis of higher demand comprehension tasks in PSA. For each cluster, we provide peak MNI coordinates, cluster size, anatomical label (defined according to the Harvard-Oxford atlas) and Brodmann Area (determined using the Talairach Daemon atlas). The ALE map was thresholded with a voxel-wise uncorrected p<0.001 cluster-forming threshold and a cluster-wise family-wise error corrected threshold of p<0.05 based on 1000 random permutations. Abbreviations: ALE = Activation Likelihood Estimation; BA = Brodmann Area; L=Left; MNI = Montreal Neurological Institute; R = Right. PSA = Person(s) with Post-stroke aphasia. * = No Brodmann Area according to Talairach Daemon.

Supplementary Table S20: Significant clusters from the ALE map of PSA performing lower demand comprehension tasks

| **Single dataset analysis** | 178 foci  12 subject groups  76 subjects | |  |  |  |  |
| --- | --- | --- | --- | --- | --- | --- |
| **Cluster** | **Cluster size (mm^3^)** | **x** | **y** | **z** | **Label** | **BA** |
| 1 | 1632 | -58 | 4 | -20 | L Temporal Pole | 21 |
|  |  | -52 | 12 | -20 | L Temporal Pole | 38 |
| 2 | 920 | -52 | -38 | -2 | L posterior Middle Temporal Gyrus | * |
|  |  | -60 | -40 | -4 | L posterior Middle Temporal Gyrus | 21 |
| 3 | 880 | -42 | 20 | -32 | L Temporal Pole | 38 |

Table of clusters produced by the ALE single dataset analysis of lower demand comprehension tasks in PSA. For each cluster, we provide peak MNI coordinates, cluster size, anatomical label (defined according to the Harvard-Oxford atlas) and Brodmann Area (determined using the Talairach Daemon atlas). The ALE map was thresholded with a voxel-wise uncorrected p<0.001 cluster-forming threshold and a cluster-wise family-wise error corrected threshold of p<0.05 based on 1000 random permutations. Abbreviations: ALE = Activation Likelihood Estimation; BA = Brodmann Area; L=Left; MNI = Montreal Neurological Institute; R = Right. PSA = Person(s) with Post-stroke aphasia. * = No Brodmann Area according to Talairach Daemon.

Supplementary Table S21: Significant clusters from the ALE map of controls performing higher demand comprehension tasks

| **Single dataset analysis** | 278 foci  18 subject groups  280 subjects | |  |  |  |  |
| --- | --- | --- | --- | --- | --- | --- |
| **Cluster** | **Cluster size (mm^3^)** | **x** | **y** | **z** | **Label** | **BA** |
| 1 | 2648 | -42 | 12 | 28 | L Inferior Frontal Gyrus pars opercularis | 9 |
|  |  | -46 | 28 | 26 | L Middle Frontal Gyrus | 9 |
|  |  | -46 | 14 | 12 | L Inferior Frontal Gyrus pars opercularis | 13 |
|  |  | -44 | 24 | 30 | L Middle Frontal Gyrus | 9 |
|  |  | -52 | 10 | 8 | L Inferior Frontal Gyrus pars opercularis | 44 |
| 2 | 1920 | -48 | -52 | -10 | L temporooccipital Inferior Temporal Gyrus | 37 |
|  |  | -46 | -50 | -16 | L temporooccipital Inferior Temporal Gyrus | 37 |
|  |  | -52 | -44 | -8 | L temporooccipital Middle Temporal Gyrus | 37 |
|  |  | -50 | -60 | -18 | L temporooccipital Inferior Temporal Gyrus | 37 |
| 3 | 1120 | -6 | 54 | 24 | Superior Frontal Gyrus | 9 |
|  |  | 0 | 44 | 24 | Paracingulate Gyrus | 9 |
|  |  | 0 | 42 | 28 | Paracingulate Gyrus | 9 |
|  |  | -4 | 46 | 36 | Superior Frontal Gyrus | 8 |
| 4 | 1016 | -6 | 22 | 48 | Paracingulate Gyrus | 8 |
|  |  | -2 | 36 | 52 | Superior Frontal Gyrus | 8 |
|  |  | -6 | 32 | 46 | Superior Frontal Gyrus | 8 |
| 5 | 816 | -38 | 30 | -8 | L Frontal Orbital Cortex | 47 |

Table of clusters produced by the ALE single dataset analysis of higher demand comprehension tasks in controls. For each cluster, we provide peak MNI coordinates, cluster size, anatomical label (defined according to the Harvard-Oxford atlas) and Brodmann Area (determined using the Talairach Daemon atlas). The ALE map was thresholded with a voxel-wise uncorrected p<0.001 cluster-forming threshold and a cluster-wise family-wise error corrected threshold of p<0.05 based on 1000 random permutations. Abbreviations: ALE = Activation Likelihood Estimation; BA = Brodmann Area; L=Left; MNI = Montreal Neurological Institute; R = Right. PSA = Person(s) with Post-stroke aphasia. * = No Brodmann Area according to Talairach Daemon.

Supplementary Table S22: Group similarities between the ALE maps of PSA and controls performing higher demand comprehension tasks

| **Conjunction** |  |  |  |  |  |  |
| --- | --- | --- | --- | --- | --- | --- |
| **Cluster** | **Cluster size (mm^3^)** | **x** | **y** | **z** | **Label** | **BA** |
| 1 | 576 | -46 | 14 | 14 | L Inferior Frontal Gyrus pars opercularis | 13 |
|  |  | -42 | 16 | 24 | L Inferior Frontal Gyrus pars opercularis | 9 |
|  |  | -46 | 14 | 24 | L Inferior Frontal Gyrus pars opercularis | 9 |
| 2 | 400 | -36 | 30 | -10 | L Frontal Orbital Cortex | 47 |
|  |  | -40 | 30 | -6 | L Frontal Orbital Cortex | 47 |
| 3 | 200 | -44 | 28 | 24 | L Middle Frontal Gyrus | 46 |
| 4 | 96 | -8 | 18 | 48 | Superior Frontal Gyrus | 32 |

Table of clusters produced by the ALE contrast analysis showing the conjunction between higher demand comprehension tasks in PSA and in controls. For each cluster, we provide peak MNI coordinates, cluster size, anatomical label (defined according to the Harvard-Oxford atlas) and Brodmann Area (determined using the Talairach Daemon atlas). Abbreviations: ALE = Activation Likelihood Estimation; BA = Brodmann Area; L=Left; MNI = Montreal Neurological Institute; R = Right. PSA = Person(s) with Post-stroke aphasia. * = No Brodmann Area according to Talairach Daemon.

Supplementary Table S23: Group differences between the ALE maps of PSA versus controls performing higher demand comprehension tasks

| **Cluster** | **Cluster size (mm^3^)** | **x** | **y** | **z** | **Z (peak)** | **Label** | **BA** |
| --- | --- | --- | --- | --- | --- | --- | --- |
| **PSA>Controls (higher demand comprehension)** |  |  |  |  |  |  |  |
| 1 | 736 | 46 | 23 | 19 | 2.23 | R Inferior Frontal Gyrus pars triangularis | 46 |
|  |  | 40 | 20 | 18 | 2.27 | R Inferior Frontal Gyrus pars opercularis | * |
|  |  | 44 | 16 | 22 | 2.18 | R Inferior Frontal Gyrus pars opercularis | 9 |
| 2 | 672 | 34 | 18 | 4 | 2.43 | R Insular Cortex | * |
| 3 | 616 | -56 | -6 | -8 | 2.71 | L anterior Superior Temporal Gyrus | 22 |
| 4 | 272 | 48 | 30 | -12 | 2.19 | R Frontal Orbital Cortex | 47 |
| **Controls>PSA (higher demand comprehension)** |  |  |  |  |  |  |  |
| 1 | 912 | -4 | 48 | 28 | 2.442152 | Superior Frontal Gyrus | 9 |
|  |  | 0 | 44 | 22 | 1.747215 | Paracingulate Gyrus | 9 |
| 2 | 336 | -44 | 16 | 32 | 2.180776 | L Middle Frontal Gyrus | 9 |
|  |  | -44 | 12 | 26 | 1.925235 | L Inferior Frontal Gyrus pars opercularis | 9 |
| 3 | 200 | -4 | 30 | 50 | 2.232226 | Superior Frontal Gyrus | 8 |

Table of clusters produced by the ALE contrast analysis showing differences between higher demand comprehension tasks in PSA vs. in controls. For each cluster, we provide the peak MNI coordinate, cluster size, anatomical label (defined according to the Harvard-Oxford atlas) and Brodmann Area (determined using the Talairach Daemon atlas). Thresholded ALE maps from the two datasets being compared were subtracted from each other and thresholded at p<0.05 using 10000 P-value permutations with a minimum cluster threshold of 200mm^3^. Abbreviations: ALE = Activation Likelihood Estimation; BA = Brodmann Area; L=Left; MNI = Montreal Neurological Institute; R = Right. PSA = Person(s) with Post-stroke aphasia. * = No Brodmann Area according to Talairach Daemon.

Supplementary Table S24: Group differences between the ALE maps of PSA performing higher demand versus lower demand comprehension tasks

| **Cluster** | **Cluster size (mm^3^)** | **x** | **y** | **Z** | **Z (peak)** | **Label** | **BA** |
| --- | --- | --- | --- | --- | --- | --- | --- |
| **Higher Demand Comprehension>Lower Demand Comprehension (PSA)** |  |  |  |  |  |  |  |
| 1 | 4904 | 40 | 24 | 2 | 3.24 | R Frontal Operculum Cortex | 13 |
|  |  | 34 | 20 | -6 | 2.85 | R Insular Cortex | * |
|  |  | 30 | 18 | 2 | 2.67 | R Insular Cortex | * |
| 2 | 3064 | -44 | 27 | 18 | 3.89 | L Inferior Frontal Gyrus pars triangularis | 46 |
|  |  | -41 | 22 | 18 | 3.72 | L Inferior Frontal Gyrus pars opercularis | 46 |
|  |  | -42 | 21 | 23 | 3.54 | L Inferior Frontal Gyrus pars opercularis | 46 |
| 3 | 1488 | -42 | -70 | 24 | 2.30 | L superior Lateral Occipital Cortex | 39 |
|  |  | -47 | -70 | 22 | 2.25 | L superior Lateral Occipital Cortex | 39 |
|  |  | -42 | -66 | 18 | 2.24 | L superior Lateral Occipital Cortex | 19 |
| 4 | 1376 | 45 | 27 | 18 | 2.69 | R Inferior Frontal Gyrus pars triangularis | 46 |
|  |  | 40 | 22 | 18 | 2.61 | R Inferior Frontal Gyrus pars triangularis | * |
| **Lower Demand Comprehension> Higher Demand Comprehension (PSA)** |  |  |  |  |  |  |  |
| 1 | 1584 | -60 | 8 | -26 | 2.45 | L Temporal Pole | 21 |
|  |  | -62 | 6 | -22 | 2.45 | L Temporal Pole | 21 |
|  |  | -56 | 9 | -23 | 2.42 | L Temporal Pole | 21 |
|  |  | -52 | 20 | -20 | 1.88 | L Temporal Pole | 38 |
| 2 | 880 | -44 | 17 | -33 | 2.15 | L Temporal Pole | 38 |
|  |  | -43 | 23 | -33 | 2.15 | L Temporal Pole | 38 |

Table of clusters produced by the ALE contrast analysis showing differences between PSA performing higher demand comprehension tasks versus PSA performing lower demand comprehension tasks. For each cluster, we provide the peak MNI coordinate, cluster size, anatomical label (defined according to the Harvard-Oxford atlas) and Brodmann Area (determined using the Talairach Daemon atlas). Thresholded ALE maps from the two datasets being compared were subtracted from each other and thresholded at p<0.05 using 10000 P-value permutations with a minimum cluster threshold of 200mm^3^. Abbreviations: ALE = Activation Likelihood Estimation; BA = Brodmann Area; L=Left; MNI = Montreal Neurological Institute; R = Right. PSA = Person(s) with Post-stroke aphasia. * = No Brodmann Area according to Talairach Daemon.

Supplementary Table S25: Significant clusters from the ALE map of PSA performing higher demand production tasks

| **Single dataset analysis** | 178 foci  13 subject groups  118 subjects | |  |  |  |  |
| --- | --- | --- | --- | --- | --- | --- |
| **Cluster** | **Cluster size (mm^3^)** | **x** | **y** | **z** | **Label** | **BA** |
| 1 | 1024 | 52 | -18 | 10 | R Heschl’s Gyrus (includes H1 and H2) | 41 |
|  |  | 56 | -26 | 4 | R posterior Superior Temporal Gyrus | 41 |
|  |  | 44 | -22 | 12 | R Heschl’s Gyrus (includes H1 and H2) | 41 |
|  |  | 50 | -28 | 4 | R posterior Superior Temporal Gyrus | 41 |
|  |  | 52 | -24 | 2 | R posterior Superior Temporal Gyrus | 41 |
| 2 | 728 | 48 | -6 | 36 | R Precentral Gyrus | 6 |
| 3 | 560 | -48 | 32 | 10 | L Inferior Frontal Gyrus pars triangularis | 46 |
|  |  | -48 | 34 | 14 | L Inferior Frontal Gyrus pars triangularis | 46 |
|  |  | -54 | 34 | 20 | L Middle Frontal Gyrus | 46 |

Table of clusters produced by the ALE single dataset analysis of higher demand production tasks in PSA. For each cluster, we provide peak MNI coordinates, cluster size, anatomical label (defined according to the Harvard-Oxford atlas) and Brodmann Area (determined using the Talairach Daemon atlas). The ALE map was thresholded with a voxel-wise uncorrected p<0.001 cluster-forming threshold and a cluster-wise family-wise error corrected threshold of p<0.05 based on 1000 random permutations. Abbreviations: ALE = Activation Likelihood Estimation; BA = Brodmann Area; L=Left; MNI = Montreal Neurological Institute; R = Right. PSA = Person(s) with Post-stroke aphasia. * = No Brodmann Area according to Talairach Daemon.

Supplementary Table S26: Significant clusters from the ALE map of PSA performing lower demand production tasks

| **Single dataset analysis** | 325 foci  29 subject groups  107 subjects | |  |  |  |  |
| --- | --- | --- | --- | --- | --- | --- |
| **Cluster** | **Cluster size (mm^3^)** | **x** | **y** | **z** | **Label** | **BA** |
| 1 | 2896 | 56 | -30 | 2 | R posterior Superior Temporal Gyrus | 22 |
|  |  | 66 | -22 | -2 | R posterior Superior Temporal Gyrus | 21 |
| 2 | 1040 | 54 | 4 | 36 | R Precentral Gyrus | 6 |
|  |  | 56 | 0 | 38 | R Precentral Gyrus | 6 |

Table of clusters produced by the ALE single dataset analysis of lower demand production tasks in PSA. For each cluster, we provide peak MNI coordinates, cluster size, anatomical label (defined according to the Harvard-Oxford atlas) and Brodmann Area (determined using the Talairach Daemon atlas). The ALE map was thresholded with a voxel-wise uncorrected p<0.001 cluster-forming threshold and a cluster-wise family-wise error corrected threshold of p<0.05 based on 1000 random permutations. Abbreviations: ALE = Activation Likelihood Estimation; BA = Brodmann Area; L=Left; MNI = Montreal Neurological Institute; R = Right. PSA = Person(s) with Post-stroke aphasia. * = No Brodmann Area according to Talairach Daemon.

Supplementary Table S27: Significant clusters from the ALE map of controls performing lower demand production tasks

| **Single dataset analysis** | 232 foci  10 subject groups  119 subjects | |  |  |  |  |
| --- | --- | --- | --- | --- | --- | --- |
| **Cluster** | **Cluster size (mm^3^)** | **x** | **y** | **z** | **Label** | **BA** |
| 1 | 1896 | -4 | 2 | 60 | Supplementary Motor Cortex | 6 |
|  |  | -4 | 12 | 52 | Paracingulate Gyrus | 6 |
|  |  | -6 | 0 | 70 | Supplementary Motor Cortex | 6 |
| 2 | 1848 | -62 | -42 | 6 | L posterior Supramarginal Gyrus | 22 |
|  |  | -52 | -34 | 8 | L Planum Temporale | 22 |

Table of clusters produced by the ALE single dataset analysis of lower demand production tasks in controls. For each cluster, we provide peak MNI coordinates, cluster size, anatomical label (defined according to the Harvard-Oxford atlas) and Brodmann Area (determined using the Talairach Daemon atlas). The ALE map was thresholded with a voxel-wise uncorrected p<0.001 cluster-forming threshold and a cluster-wise family-wise error corrected threshold of p<0.05 based on 1000 random permutations. Abbreviations: ALE = Activation Likelihood Estimation; BA = Brodmann Area; L=Left; MNI = Montreal Neurological Institute; R = Right. PSA = Person(s) with Post-stroke aphasia. * = No Brodmann Area according to Talairach Daemon.

Supplementary Table S28: Group similarities between the ALE maps of PSA and controls performing lower demand production tasks

| **Conjunction** |  |  |  |  |  |  |
| --- | --- | --- | --- | --- | --- | --- |
| **Cluster** | **Cluster size (mm^3^)** | **x** | **y** | **z** | **Label** | **BA** |
| No clusters found |  |  |  |  |  |  |

Table of clusters produced by the ALE contrast analysis showing the conjunction between lower demand production tasks in PSA and in controls. For each cluster, we provide peak MNI coordinates, cluster size, anatomical label (defined according to the Harvard-Oxford atlas) and Brodmann Area (determined using the Talairach Daemon atlas). Abbreviations: ALE = Activation Likelihood Estimation; BA = Brodmann Area; L=Left; MNI = Montreal Neurological Institute; R = Right. PSA = Person(s) with Post-stroke aphasia. * = No Brodmann Area according to Talairach Daemon.

Supplementary Table S29: Group differences between the ALE maps of PSA versus controls performing lower demand production tasks

| **Cluster** | **Cluster size (mm^3^)** | **x** | **y** | **z** | **Z (peak)** | **Label** | **BA** |
| --- | --- | --- | --- | --- | --- | --- | --- |
| **PSA>Controls (lower demand production)** |  |  |  |  |  |  |  |
| No clusters found |  |  |  |  |  |  |  |
| **Controls>PSA (lower demand production)** |  |  |  |  |  |  |  |
| 1 | 2112 | -3 | 8 | 54 | 3.35 | Supplementary Motor Cortex | 6 |
| 2 | 2032 | -59 | -40 | 8 | 3.89 | L posterior Superior Temporal Gyrus | 22 |
|  |  | -48 | -32 | 10 | 3.16 | L Planum Temporale | 41 |
|  |  | -56 | -36 | 8 | 2.82 | L posterior Superior Temporal Gyrus | 22 |
| 3 | 352 | 64 | -32 | 6 | 2.07 | R posterior Superior Temporal Gyrus | 42 |
|  |  | 64 | -26 | 4 | 2.05 | R posterior Superior Temporal Gyrus | 22 |
| 4 | 224 | 48 | -4 | 34 | 2.31 | R Precentral Gyrus | 6 |

Table of clusters produced by the ALE contrast analysis showing differences between lower demand production tasks in PSA vs. in controls. Thresholded ALE maps from the two datasets being compared were subtracted from each other and thresholded at p<0.05 using 10000 P-value permutations with a minimum cluster threshold of 200mm^3^. Abbreviations: ALE = Activation Likelihood Estimation; BA = Brodmann Area; L=Left; MNI = Montreal Neurological Institute; R = Right. PSA = Person(s) with Post-stroke aphasia. * = No Brodmann Area according to Talairach Daemon.

Supplementary Table S30: Group differences between the ALE maps of PSA performing higher demand versus lower demand production tasks

| **Cluster** | **Cluster size (mm^3^)** | **x** | **y** | **z** | **Z (peak)** | **Label** | **BA** |
| --- | --- | --- | --- | --- | --- | --- | --- |
| **Higher Demand Production> Lower Demand Production (PSA)** |  |  |  |  |  |  |  |
| 1 | 416 | 48 | -6 | 40 | 2.09 | R Precentral Gyrus | 6 |
| 2 | 328 | 50 | -22 | 12 | 1.95 | R Heschl’s Gyrus (includes H1 and H2) | 41 |
|  |  | 56 | -22 | 10 | 1.93 | R Planum Temporale | 41 |
| 3 | 288 | 52 | 16 | 0 | 2.16 | R Inferior Frontal Gyrus pars opercularis | 13 |
|  |  | 48 | 18 | 2 | 2.08 | R Frontal Operculum Cortex | 44 |
| **Lower Demand Production> Higher Demand Production (PSA)** |  |  |  |  |  |  |  |
| No clusters found |  |  |  |  |  |  |  |

Table of clusters produced by the ALE contrast analysis showing differences between PSA performing higher demand production tasks versus PSA performing lower demand production tasks. For each cluster, we provide the peak MNI coordinate, cluster size, anatomical label (defined according to the Harvard-Oxford atlas) and Brodmann Area (determined using the Talairach Daemon atlas). Thresholded ALE maps from the two datasets being compared were subtracted from each other and thresholded at p<0.05 using 10000 P-value permutations with a minimum cluster threshold of 200mm^3^. Abbreviations: ALE = Activation Likelihood Estimation; BA = Brodmann Area; L=Left; MNI = Montreal Neurological Institute; R = Right. PSA = Person(s) with Post-stroke aphasia. * = No Brodmann Area according to Talairach Daemon.

Supplementary Table S31: Mean ages of the subject groups contributing to contrast ALE meta-analyses in this paper

| **Dataset 1** | **Mean ages of subject groups in Dataset 1 (median, IQR)** | **Dataset 2** | **Mean ages of subject groups in Dataset 2 (median, IQR)** | **P value** |
| --- | --- | --- | --- | --- |
| Controls, all language tasks | 57.0 (8.2) | PSA, all language tasks | 57.4 (9.0) | 0.18 |
| Controls, comprehension tasks | 57.0 (6.9) | PSA, comprehension tasks | 57.0 (9.5) | 0.48 |
| Controls, production tasks | 54.5 (8.2) | PSA, production tasks | 57.4 (10.3) | 0.14 |
| Controls, comprehension tasks | 57.0 (6.9) | Controls, production tasks | 54.5 (8.2) | 0.33 |
| PSA, comprehension tasks | 57.0 (9.5) | PSA, production tasks | 57.4 (10.3) | 0.91 |
| PSA, higher demand comprehension tasks | 58.0 (9.8) | Controls, higher demand comprehension tasks | 57.0 (5.7) | 0.53 |
| PSA, higher demand comprehension tasks | 58.0 (9.8) | PSA, lower demand comprehension tasks | 56.3 (11.1) | 0.44 |
| PSA, lower demand production tasks | 57.0 (9.1) | Controls, lower demand production tasks | 56.8 (7.0) | 0.70 |
| PSA, higher demand production tasks | 58.1 (13.4) | PSA, lower demand production tasks | 57.0 (9.1) | 0.59 |

This table compares the mean ages of subject groups in each pair of datasets for every contrast ALE meta-analysis in this paper. ‘P value’ corresponds to uncorrected two-sided p-values from Mann-Whitney U-tests comparing the mean ages of subject groups in Dataset 1 to the mean ages of subject groups in Dataset 2. Abbreviations: IQR = Interquartile Range.
